# Multidrug resistance plasmids underlie clonal expansions and international spread of *Salmonella enterica* serotype 4,[5],12,i:- ST34 in Southeast Asia

**DOI:** 10.1101/2023.01.10.523379

**Authors:** Hao Chung The, Phuong Pham, Ha Thanh Tuyen, Linh Vo Kim Phuong, Nguyen Phuong Yen, Son-Nam H Le, Duong Vu Thuy, Tran Thi Hong Chau, Hoang Le Phuc, Nguyen Minh Ngoc, Lu Lan Vi, Alison E. Mather, Guy E. Thwaites, Nicholas R. Thomson, Stephen Baker, Duy Thanh Pham

**Affiliations:** Oxford University Clinical Research Unit, Ho Chi Minh City, Vietnam; School of Biotechnology, International University, Vietnam National University, Ho Chi Minh City, Vietnam; Children’s Hospital No. 1, Ho Chi Minh City, Vietnam; Children’s Hospital No. 2, Ho Chi Minh City, Vietnam; The Hospital for Tropical Diseases, Ho Chi Minh City, Vietnam; Quadram Institute Bioscience, Norwich Research Park, Norwich, United Kingdom; University of East Anglia, Norwich, United Kingdom; Centre for Tropical Medicine and Global Health, Nuffield Department of Clinical Medicine, University of Oxford, Oxford, United Kingdom; London School of Hygiene and Tropical Medicine, London, United Kingdom; The Wellcome Sanger Institute, Hinxton, Cambridge, United Kingdom; Department of Medicine, Cambridge Institute of Therapeutic Immunology and Infectious Diseases (CITIID), University of Cambridge, Cambridge, United Kingdom

**Author notes:** Corresponding author: Dr. Hao Chung The, Department of Molecular Epidemiology, Oxford University Clinical Research Unit (OUCRU), 764 Vo Van Kiet St., Ward 1, District 5, Ho Chi Minh City, Vietnam.

## Abstract

*Salmonella enterica* serotype 4,[5],12,i:- (Typhimurium monophasic variant) of sequence type (ST) 34 has emerged as the predominant pandemic genotype in recent decades. Despite increasing reports of resistance to antimicrobials in Southeast Asia, *Salmonella* ST34 population structure and evolution remained understudied in the region. Here we performed detailed genomic investigations on 454 ST34 genomes collected from Vietnam and diverse geographical sources to elucidate the pathogen’s epidemiology, evolution and antimicrobial resistance. We showed that ST34 has been introduced into Vietnam in at least nine occasions since 2000, forming five co-circulating major clones responsible for paediatric diarrhoea and bloodstream infection. Most expansion events were associated with acquisitions of large multidrug resistance plasmids of IncHI2 or IncA/C2. Particularly, the self-conjugative IncA/C2 pST34VN2 (co-transferring *bla*_CTX-M-55_, *mcr-3.1*, and *qnrS1*) underlies local expansion and intercontinental spread in two separate ST34 clones. At the global scale, Southeast Asia was identified as a hotspot for the emergence and dissemination of multidrug resistant *Salmonella* ST34, and mutation analysis suggests of selection in antimicrobial responses and key virulence factors. Our work enriches the understanding on epidemiology and evolution of this variant in Southeast Asia, and determines that multidrug resistance plasmids have driven its local and potentially global success.

## Introduction

Nontyphoidal *Salmonella enterica* (NTS) rank among the most common bacterial pathogens causing diarrheal diseases worldwide, leading to an estimate of 75 million cases per year^1^. NTS can also cause highly fatal invasive diseases in vulnerable populations, such as young children and HIV-positive patients in Asia and Africa, with nearly 77,000 attributed deaths globally ^2^. There are >2,500 recorded NTS serotypes and their distributions vary geographically and temporally, but few stand out to dominate the global epidemiology. Chief among these is *S*. Typhimurium, a host generalist capable of surviving in a broad range of animals (mainly poultry, swine, and cattle), human and the environment. This serotype is further delineated into several related sequence types (STs), with ST19, ST36, ST313 and ST34 attributing to the majority of global disease burden ^3,4^. In addition, serotype variability also arises due to losses or inactivations of the gene encoding the antigenic phase II flagellin (*fljB*), forming the monophasic Typhimurium variant (serotype *S*. 4,[5],12,i:-). In the last decade, *S*. 4,[5],12,i:-, most frequently ascribed to ST34, has emerged as an important pandemic variant with increasing antimicrobial resistance (AMR) ^5–7^. This variant is the culprit of multiple foodborne outbreaks, and recently responsible for a multi-country outbreak linked to chocolate products in Europe, triggering large-scale product recalls and substantial economic loss ^8^.

The evolution and epidemiology of ST34 has been investigated extensively in high income settings, including the UK, USA, Japan and Australia ^9–12^. These showed that ST34 has likely diverged from the ancestral ST19 in the mid 1990s ^10,13^, and it is characterized by several genetic features: (a) frequent deletion(s) of *fljB* and the IncFII *S*. Typhimurium virulence plasmid, (b) acquisition of the genomic island SGI-4 enhancing resistance to copper ^14^, and (c) chromosomal integration of AMR genes conferring the ASSuT resistance pattern (to ampicillin, streptomycin, sulphonamides and tetracycline) ^10^. Recently, ST34 harbouring extensive spectrum beta-lactamases (ESBLs) or mobile colistin resistance determinants *mcr* have been reported in China and several Southeast Asia countries ^15–17^. Resistance to these critically important antimicrobials (CIAs) sparked great concern as they are frequently prescribed for treatment of gastroenteritis and severe infections ^18^. Besides, these mobile AMR elements in *Salmonella* could act as reservoir for further dissemination to commensal and pathogenic enteric bacteria.

Vietnam currently ranks among the top ten producers in the global pork industry, and the country was estimated to have the highest prevalence of AMR *S*. Typhimurium in Southeast Asia^19^. Our previous work has shown that NTS gradually became the predominant bacterial aetiology of dysentery ^20^, and *S*. 4,[5],12,i:-ST34 accounted for nearly one third of this NTS burden ^21^. Furthermore, the majority of collected ST34 were multidrug resistant (MDR), with frequent resistance to ceftriaxone and azithromycin. Given that ST34 genomic epidemiology remains relatively unexplored in Southeast Asia, this study aims to use whole genome analyses to unravel the population structure of Vietnamese ST34 in regional and global phylogenetic context, as well as to understand the basis of its MDR genotype.

## Results

### Vietnamese *Salmonella enterica* ST34 in global context

We utilized a collection of 133 *S. enterica* ST34 genome sequences, derived from a diarrheal surveillance study conducted in Southern Vietnam from 2014 to 2016 (Table 1). To provide phylogenetic context for our investigation, we gathered a collection of contemporary ST34 sequences originating from Vietnam (n=77 isolated from 2007 to 2015; previously published) ^22^ and other countries (n=244) (Table 1). Since there is limited information on the genomic epidemiology of ST34 in Asia, we preferentially selected genomes originating from this region. This resulted in an over-representation of Asian ST34 (315/454; from Vietnam, Thailand, China, Japan, Taiwan, Cambodia and Laos), spanning a period of 18 years (2002 - 2019). The majority of isolates (332/454) originated from humans, while animal isolates (118/454) included those from swine, cattle, poultry and fish. For Vietnamese human-derived isolates, of which clinical data were available, gastroenteritis was the most common manifestation (n=150), followed by bloodstream infections (n=26). Details of each isolate are provided in Table S1.

**Table 1.** Summary of *Salmonella enterica* ST34 genomes used in this study.

| Region | Country | Reference | No. | Duration |
| --- | --- | --- | --- | --- |
| Southeast Asia | Vietnam | Duong et al., JCM, 2020 | 133 | 2014 – 2016 |
|  | Vietnam | Mather et al., mBio, 2018 | 77 | 2007 – 2015 |
|  | Cambodia | Enterobase | 4 | 2016 – 2018 |
|  | Thailand | Enterobase | 15 | 2009 - 2019 |
|  | Laos | Enterobase | 4 |  |
|  | Unknown | Ingle et al., Nat Comms, 2021 | 40 | 2007 – 2016 |
| East Asia | China | Enterobase | 14 | 2012 – 2018 |
|  | Japan | Enterobase | 13 | 2002 – 2017 |
|  | Taiwan | Enterobase | 15 | 2012 – 2018 |
| Oceania | Australia | Ingle et al., Nat Comms, 2021 | 38 | 2007 – 2017 |
| Europe | UK | Petrovska et al., EID, 2016<br>Arnott et al., EID, 2018 | 56 | 2006 – 2017 |
|  | Italy | Petrovska et al., EID, 2016 | 13 | 2006 – 2010 |
| America | USA | Elnekave et al., EID, 2020 | 32 | 2011 – 2016 |
|  |  |  | 454 |  |

Hierarchical Bayesian clustering on the genetic variation of 454 ST34 genomes delineated them into four lineages (named herein as BAPS-1 to -4), which agreed with its phylogenetic groupings on the maximum likelihood (ML) phylogeny (Figure 1). Molecular serotyping indicated that the serotype 4,[5],12:i:-constituted ~87% in our collection (n=397/454). This serotype made up 62% of the ancestral lineage BAPS-1 and >98% of two expansive lineages (BAPS-3 and -4). The remaining lineage (BAPS-2) was exclusively composed of Vietnamese isolates and mostly of serotype Typhimurium (n=30/32), similar to our previous report ^22^. Temporal phylogenetic inference, as implemented in BEAST v1.8.2, estimated that ST34 evolved with a substitution rate of 4.99E-7 substitutions per site per year (95% highest posterior density [HPD]: 4.27E-7 to 5.76E-7). The most recent common ancestor (MRCA) of all examined ST34 likely emerged in 1995 (95% HPD: 1992 - 1998) (Figure 2A). These estimates mirror similar calculations from recent large-scale studies of *S*. 4,[5],12:i:-ST34 ^10,23^.

**Figure 1.**
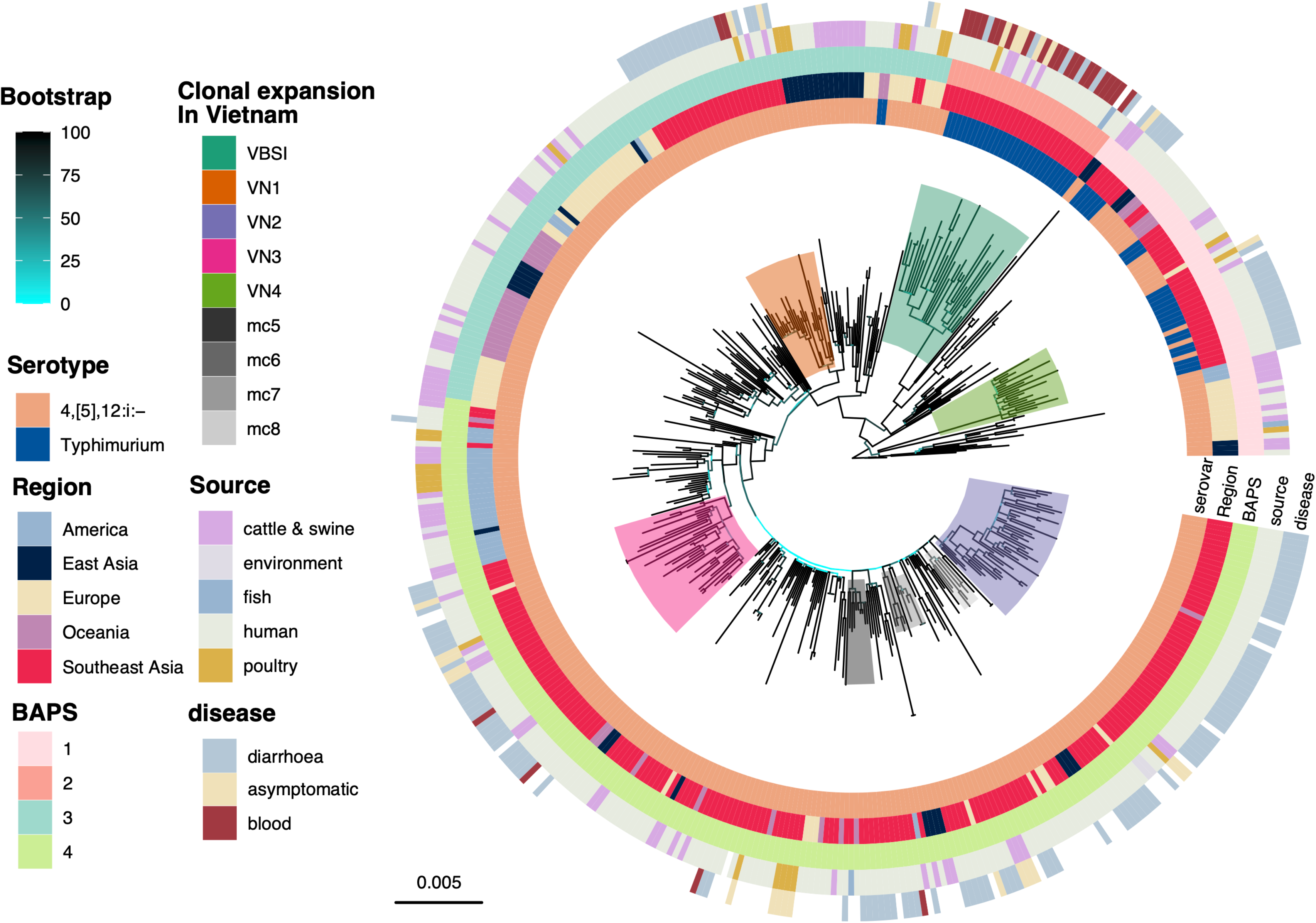
Global phylogeny of *Salmonella enterica* ST34. The figure displays the maximum likelihood phylogeny of 454 *S. enterica* ST34 isolates, constructed from 4,962 single nucleotide polymorphisms. The phylogeny is rooted using an *S. enterica* ST19 outgroup. Branches are coloured in accordance to bootstrapping values, from low (cyan) to high (black). The rings present information associated with each taxon, in the following order (inner to outermost): (1) serotype predicted *in silico* from ST34 assemblies, (2) region of origin (for travel-associated isolates, the known travel destination was recorded as the region of origin), (3) population structure determined by hierarchical Bayesian clustering (BAPS), (4) source of isolation, and (5) disease manifestation (for isolates from Vietnam with clinical data). Colour shadings denote different clades (n≥5 isolates) with clonal expansion in Vietnam, of which the majority of isolates originating from Vietnam or Southeast Asia. The horizontal bar indicates the number of substitution per site.

**Figure 2.**
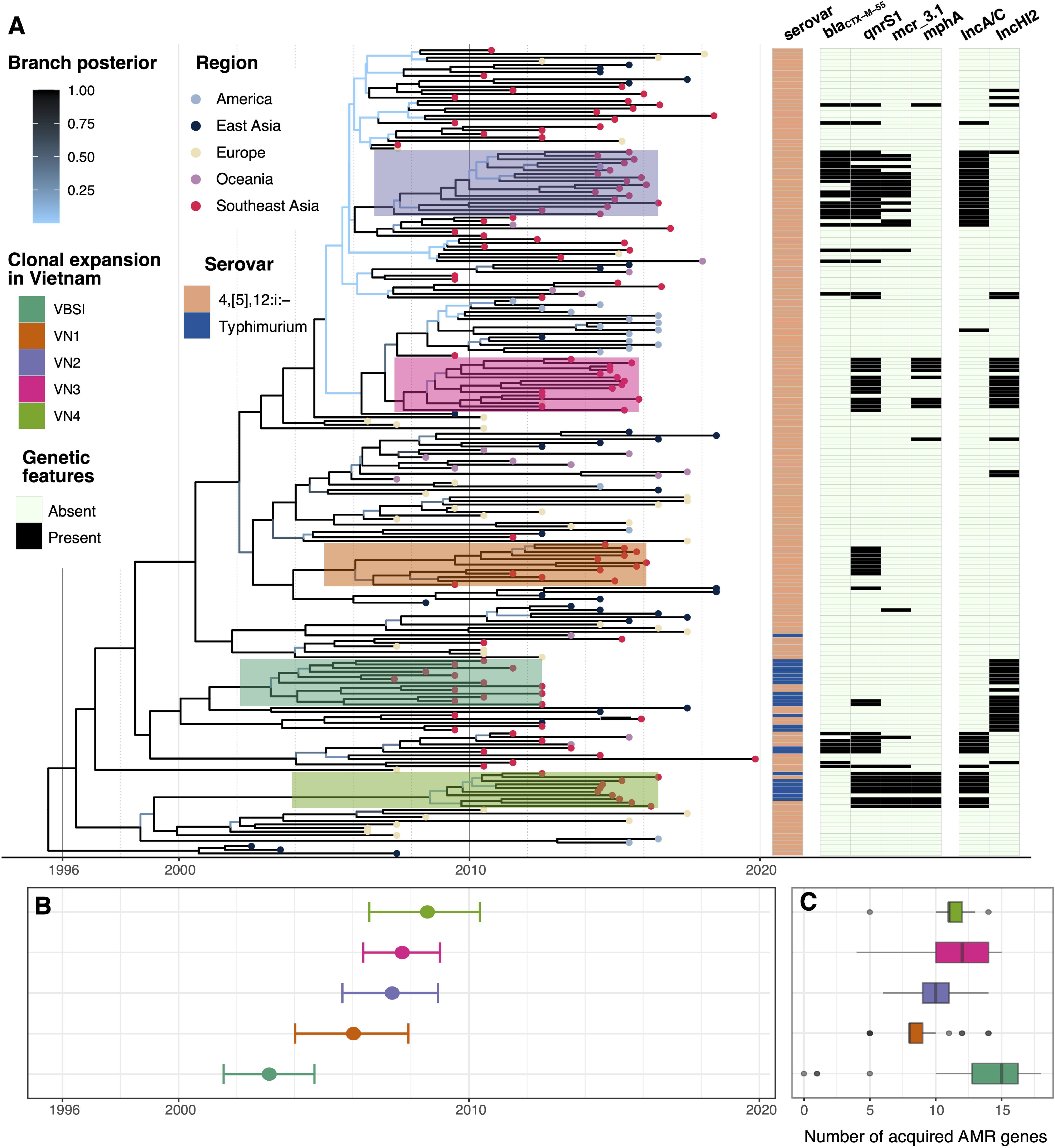
Temporal phylogenetic reconstruction of *Salmonella enterica* ST34. The panel (A) shows the maximum clade credibility (MCC) phylogeny of 222 representative *S. enterica* ST34, constructed from 2,671 single nucleotide polymorphisms. Branches are coloured according to the calculated posterior probability, from low (light blue) to high (black), while tip points are coloured based on the region of origin. The five clones associated with major clonal expansion events in Vietnam (VN1-4, VBSI) are annotated. The appended heatmap displays data associated with each taxon, including the predicted serotype (biphasic or monophasic Typhimurium), and the presence of antimicrobial resistance genes (*bla*_CTX-M-55_, *qnrS1, mcr-3.1, mphA*) and predominant plasmid replicon (IncA/C, IncHI2). (B) Estimation of the time to most common recent ancestor (tMRCA) of five major clonal expansions of ST34 (VN1-4, VBSI) in Vietnam, with bars representing the highest posterior density (HPD), and (C) Boxplots displaying the distribution (median and inter-quartile range) of acquired antimicrobial resistance genes in five major clonal expansions in Vietnam.

Though Southeast Asian isolates were distributed in all four lineages, they were disproportionally placed within lineage BAPS-4 (n=174/273), which emerged circa 2004 (95% HPD: 2003 – 2006) (Figure 1). Across the ST34 phylogeny, we identified five major clonal expansion events in Vietnam, defined as phylogenetic clusters with ≥ 90 bootstrap support and consisted of mostly Vietnamese isolates (n ≥ 20 for each clone). Four of these were associated with childhood gastroenteritis, and were named in accordance to their inferred time of divergence, including VN1 (~2006), VN2, VN3 (both ~2007) and VN4 (~2008) (Figure 2B). The remaining clone (VBSI/BAPS-2) likely arose earlier in 2003 (95% HPD: 2001 – 2004), and this clone was previously shown to cause bloodstream infections in HIV-positive patients (n=20/32) ^22^. Additionally, ST34 isolated from Vietnam were also found in several small independent clusters (mc5, mc6, mc7, mc8; each with 5 - 10 isolates), all belonging to BAPS-4. These results demonstrated that ST34 might have been introduced into Vietnam in at least nine occasions since the early 2000s, and the circulating ST34 were derived from a diverse phylogenetic background, with BAPS-4 dominating the epidemiological landscape of gastroenteritis in Vietnam and Southeast Asia.

### Southeast Asia as a source for global dissemination of *Salmonella enterica* ST34

We next sought to understand the role of Southeast Asia in the global propagation of *S*. enterica ST34. To account for the over-representation of Southeast Asian genomes in our collection, we subsampled the phylogeny to 1,000 subtrees, with each containing equal number of genomes from each geographical region (n=30). Results from stochastic mapping showed that Southeast Asia, aside from Europe, acted as the major reservoir for disseminations to America, East Asia, Europe, and Oceania on an average of 2.23 (IQR: 2 – 3), 6.02 (IRQ: 5 – 7), 5.12 (IQR: 4 – 6), and 7.92 (IQR: 7 – 9) independent events per tree, respectively (Figure 3A). Similarly, the analysis predicted that approximately 30% (IQR: 28.5% - 32.6%) of inferred evolutionary time across the ST34 phylogeny was in Southeast Asia (Figure 3B), higher than that quantified for Europe. On the other hand, Europe was inferred as a sole source of ST34 introduction into Southeast Asia, on an average of 2.95 (IQR: 2 – 3.13) independent events per tree. Conversely, analyses of ten sets of tip-location randomized phylogenies produced results that were significantly different from that estimated from the original phylogeny, ensuring that the reported inferences derived from inherent phylogeographic signal in the data (Figure S1). Particularly, the number of transition events from Southeast Asia to Oceania was higher in the true phylogeny, compared to all randomization sets (p < 0.05 in 9/10 comparisons, ANOVA-Tukey’s test [F value = 20.76, df = 10]). This evidence demonstrated that Southeast Asia acted as a major source for inter-continental transmissions of ST34.

**Figure 3.**
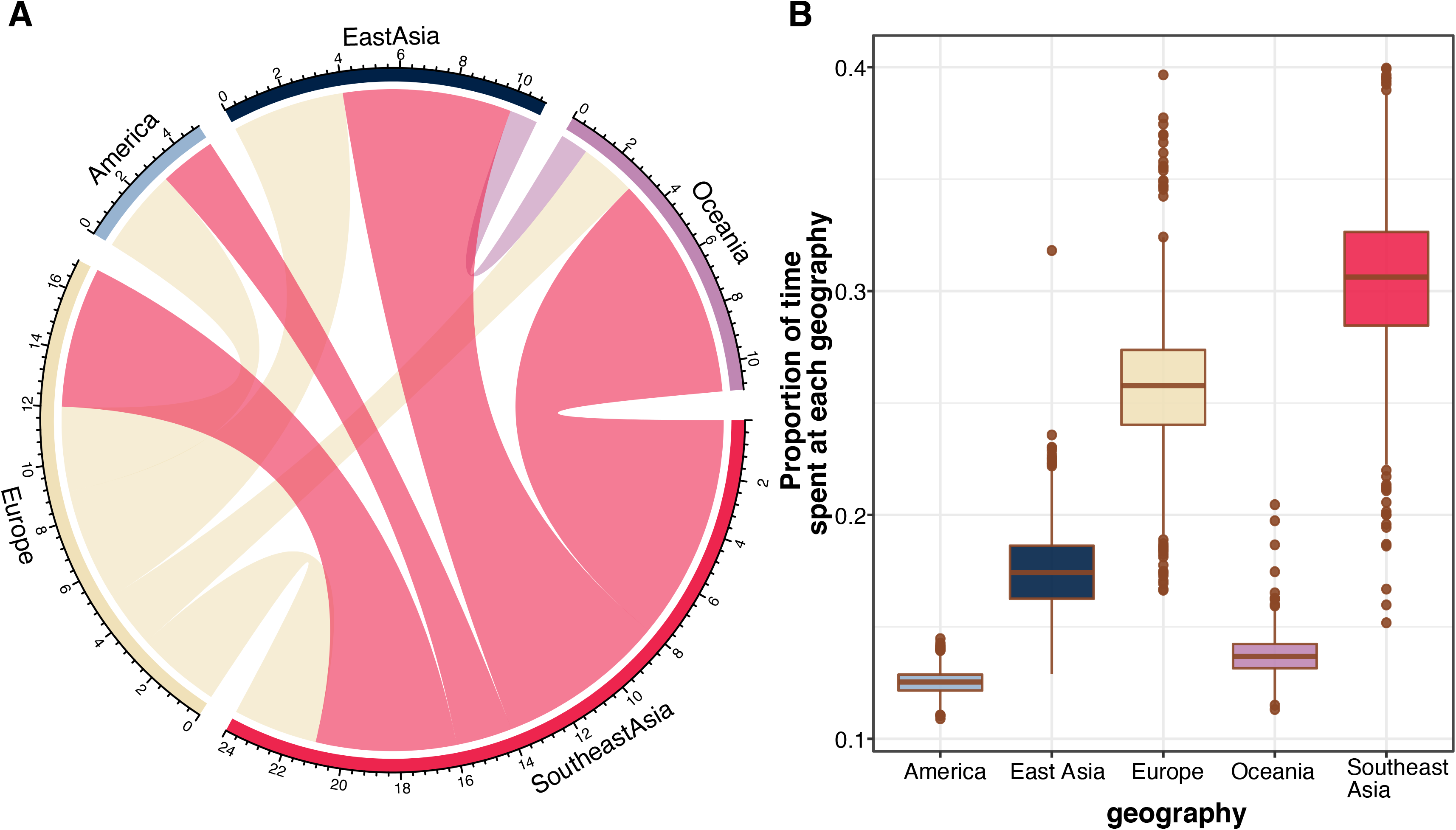
Phylogeography analysis of *Salmonella enterica* ST34. The figures display the results inferred from stochastic mapping (A) Circos plot denoting the transitions between the geographies (America, East Asia, Europe, Oceania, Southeast Asia). The broken outer ring represents the geographical sources of these transitions, proportional to their contributions to the total number of inferred transitions. Each block represents a transition direction between geographical states, with size proportional to the mean number of inferred transition events and colour based on the source of the transition. (B) The proportion of time spent in each geographical state. For both panels, the results are summarized from stochastic mapping runs of 1,000 subsamplings of the maximum likelihood phylogeny described in Figure 1.

### Mutation analysis of *Salmonella enterica* ST34

In order to gain more insights into the evolutionary trajectory of ST34, we next searched for signals of convergent evolution in the examined genomes, using annotations generated from high-quality reference-based short read mapping. Genes bearing recurring mutations among phylogenetically diverse organisms are indicators of convergent evolution, and ones with excessive amounts of nonsynonymous mutations (high dN/dS) point to evidence of positive selection. 2,310 homologues were shown to have at least one substitution during ST34 evolution, and our analyses determined that 17 genes were predicted to have undergone positive selection (adjusted dN/dS ranging from 2.07 to 3.31) (Table 2). These include genes involved in cellular response to antimicrobials (*ramR, smvA, arnA*), biosynthesis of cellular metabolite and components (*stiC, glf, cadC, bcsB, ccmA*), and response to cellular stress (*rpoS*). Particularly, RamR acts as a main transcriptional repressor of the MDR efflux pump *acrAB-tolC* complex^24^, while ArnA catalyzes the maturation of lipid A and resistance to polymyxins^25^. Notably, the remaining H antigen of the monophasic serotype (*fliC*) has the highest dN/dS estimate (= 3.31). Aside from its structural role as the flagellar monomer, FliC is also highly immunogenic and activates TLR5 responses in cells attacked by bacteria^26^. Additionally, three identified genes (*phoQ, misL, barA*) have been shown to influence *Salmonella* virulence in experimental models (Table 2). The canonical PhoPQ two-component system governs the expression of more than 60 *Salmonella* virulence genes^27^, while the BarA/SirA system positively regulates bacterial invasion via activation of SPI-1^28^. The autotransporter MisL positively impacts *Salmonella* biofilm formation and intestinal colonization in mice^29^. On the contrary, only one gene (*bapA*) was shown to be under negative selection, since no substitutions were observed in this gene, despite its great length (3,825 amino acids). Overexpression of *bapA* was shown to increase biofilm formation, and the secreted BapA contributes to invasion through oral *Salmonella* ingestion^30^.

**Table 2.** Genes identified to be under positive selection in *Salmonella enterica* ST34.

| Gene | No. of N<sup>a</sup> SNPs | No. of S<sup>b</sup> SNPs | Adjusted dN/dS | Product | GO<sup>c</sup> process |
| --- | --- | --- | --- | --- | --- |
| *stiC* | 6 | 1 | 2.48 | fimbrial outer membrane usher protein | pilus assembly |
| *fliC* | 8 | 1 | 3.31 | flagellin | Motility |
| *ramR* | 9 | 0 | Inf | Transcriptional regulator RamR | Response to antibiotic |
| *smvA* | 5 | 0 | Inf | efflux pump for methyl viologen, acriflavine and quarternary ammonium compounds | Response to antibiotic |
| *arnA* | 5 | 1 | 2.07 | bifunctional UDP-glucuronic acid oxidase/UDP-4-amino-4-deoxy-L-arabinose formyltransferase | Response to antibiotic (polymixin resistance) |
| *glf* | 5 | 1 | 2.07 | UDP-galactopyranose mutase | cell wall polysaccharide biosynthesis |
| *phoQ* | 8 | 0 | Inf | virulence sensor histidine kinase | major regulator of virulence in *Salmonella* |
| *misL* | 5 | 1 | 2.07 | autotransporter important for colonization factor | virulence |
| *sstT* | 6 | 0 | Inf | serine/threonine transporter SstT | amino acid transport |
| *ushA* | 5 | 0 | Inf | bifunctional UDP-sugar hydrolase/5'-nucleotidase UshA | nucleotide catabolism |
| *B0X74_06215* | 5 | 0 | Inf | acyl-CoA dehydrogenase |  |
| *ydhK* | 6 | 1 | 2.48 | FUSC family transporter YdhK | Unknown |
| *cadC* | 5 | 1 | 2.07 | transcriptional regulator CadC (regulate cadaverine synthesis and excretion) | transcription regulation |
| *rpoS* | 8 | 0 | Inf | RNA polymerase sigma factor RpoS | response to stress |
| *barA* | 5 | 1 | 2.07 | two-component sensor histidine kinase BarA | virulence |
| *bcsB* | 5 | 1 | 2.07 | cyclic di-GMP-binding protein | cellulose biosynthesis |
| *ccmA* | 5 | 0 | Inf | cytochrome c biogenesis ATP-binding export protein CcmA | cytochrome c biosynthesis |

<sup>a</sup> Non-synonymous single-nucleotide polymorphisms
<sup>b</sup> Synonymous single-nucleotide polymorphisms
<sup>c</sup> Gene ontology biological process

Additionally, we examined ST34 genomes for presence of pseudogenization events, and found that three genes had experienced gene loss or nonsense or frameshift mutations in more than four independent occasions. These consisted of *rpoS* (22 occasions), *btuB* (13 occasions), and *cspC* (12 occasions) (Table S2). *rpoS* frequently acquires inactivating and nonsynonymous mutations during long-term storage of isolates, so these mutations were likely laboratory artefacts and not reflective of evolution processes^31^. *btuB* encodes an outer membrane protein, importing vitamin B12 as well as cytotoxic colicins released by Gram-negative competitors^32^. Thus, disruption in *btuB* is proposed to render protection from colicins, offering higher survivability in environments both inside and outside hosts. The RNA chaperon CspC plays an indispensable role in activating the master virulence regulon PhoQP inside macrophages, and *cspC* mutants showed reduced virulence expression and invasiveness in infected mice^33^. We found that most pseudogenization events were transient and not maintained in clonal expansions, except for two lineage defining mutations. These includes frameshift mutations in (1) C839CT leading to an elongation of *sseA* encoding 3-mercaptopyruvate sulfurtransferase (inherited in all 232 BAPS-4 genomes), and (2) T452TGA leading to a shortened *pduT* encoding 1,2-propanediol utilization protein (inherited in all 32 BAPS-2/VBSI genomes) ^22^.

### Maintenance of IncA/C2 or IncHI2 multidrug resistance plasmids underlies *Salmonella enterica* ST34 clonal expansion in Vietnam

Since AMR genotypes and phenotypes were largely in high agreement in *Salmonella* ^21^, we used AMR genotyping results to document ST34’s AMR evolution. Similar to previous reports ^10,13,23^, carriage of *bla*_TEM-1_, *strAB, sul2*, and *tet* were nearly ubiquitous (ranging from 80% to 92.7%), resulting in the distinct ASSuT resistance phenotype in the majority of recorded ST34. The AMR pattern of BAPS-2/VBSI was markedly distinct from the remaining three lineages, where ~80% of isolates carried multiple aminoglycoside resistance genes (Figure 4A). On the other hand, lineages BAPS-1 and 4 displayed similar AMR profiles, with the prominent prevalence of *qnrS1, bla*_CTX-M-55_, *mph*(A), and *mcr*-3.1 (Figure 4B). These confer nonsusceptibility against critically important antimicrobials (CIAs; quinolone, 3^rd^ generation cephalosporins, macrolides, and colistin respectively)^18^. Upon examining the presence of CIA resistance elements on the ST34 phylogeny, we found that they were significantly more abundant in Southeast Asian isolates (p value < 0.001, Chi-Squared test; Figure 2A). Noticeably, the four major ST34 clones in Vietnam (VN1 - 4) each carried at least one CIA resistance genes (Figure S2), while the minor clones (mc5 – 8) were generally devoid of all these four AMR elements (n=28/29). Particularly, three CIA resistance genes were present in the majority of VN2 (*qnrS, bla*_CTX-M-55_, *mcr*3.1) and VN4 (*qnrS, mph*(A), *mcr3*.1) genomes. The four major clones carried comparative amount of acquired AMR genes (median: 8 – 12), significantly lower than that of the VBSI clone (p < 0.05, ANOVA-Tukey test) (Figure 2C). *In silico* plasmid profiling revealed that each clone harboured a distinct replicon, including p0111 (VN1), IncHI2 (VN3 and VBSI), and IncA/C2 (VN2 and VN4). We combined short and long read sequencing data from representative isolates of the two latter replicons to fully resolve these plasmid structures.

**Figure 4.**
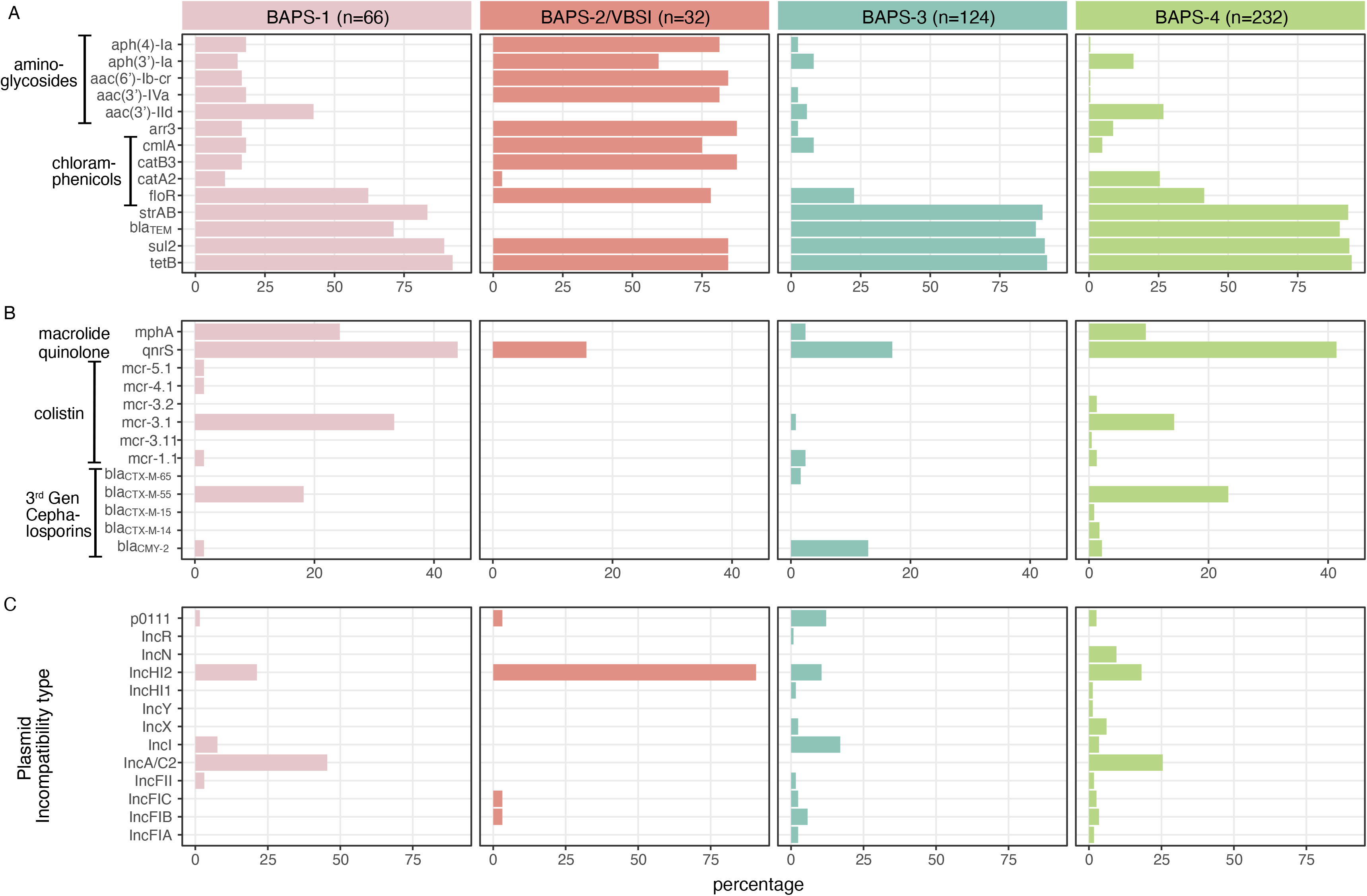
Distribution of antimicrobial resistance (AMR) genes and plasmid types among *Salmonella enterica* ST34 lineages. For each panel, the bar graph displays the percentage of isolates from each lineage (BAPS-1 to -4) carrying a respective element, stratified by (A) genes conferring resistance to aminoglycosides, rifamycin (*arr3*), chloramphenicol, ampicillin (*bla*_TEM_), sulfonamides (*sul2*), tetracycline (*tetB*), (B) genes conferring resistance to macrolides, quinolone, colistin, and 3^rd^ generation cephalosporins, and (C) plasmid incompatibility types.

Plasmid analyses confirmed that almost all AMR genes found in these major clones were co-transferred on a single MDR plasmid (IncHI2 or IncA/C2) (Figure 2A). The IncHI2 plasmids, isolated from VN3 (pST34VN3) and VBSI (pVNB151)^22^, were similar in size (>240 kbp) and each carried ~15-16 AMR genes (Table 3). Plasmid phylogeny of IncHI2 indicated that they were genetically distinct, and were respectively classified as ST2 and ST3 by pMLST scheme (Figure S3). As aforementioned, there is an excess of aminoglycoside resistance genes found in pVNB151 (*aph(3’)-Ia, aadA, aac(6’)-Iaa, aph(4)-Ia, aac(3)-IVa, aac(6’)-lb-cr*), with the latter also conferring resistance to quinolone. On the other hand, *qnrS1* and *mph(A*) were co-transferred on pST34VN3, alongside AMR determinants to phenicols (*floR, cmlA, catA2*), co-trimoxazole (*sul2-dfrA14*), ampicillin (*bla*_TEM_), lincosamide (*lnuF*) and rifampin (*arr-2*) (Table 3; Figure S4). We screened for the presence of IncHI2 plasmids in contemporary *Salmonella* genomes of other serotypes isolated in Vietnam (n=317), and found that IncHI2 was detected in several other serotypes, most frequently *S*. Newport (n=14) and *S*. Stanley (n=13) (Figure S3). Notably, pST34VN3 shared minute genetic divergence from plasmids derived from *S*. Newport ST46 (n=11), *S*. Kentucky ST198 (n=2), *S*. Corvallis and *S*. Wandsworth (n=1), among which eight was solely attributed with carriage of all three CIA resistance genes (*bla*_CTX-M-55_, *qnrS1*, and *mphA*). This implies that the IncHI2 plasmid is widely mobilized in different *Salmonella* genetic backgrounds, and recombination of AMR genes gives rise to clones with increasing resistance to CIAs.

**Table 3.** Plasmids with complete structure investigated in this study.

| Plasmid name | Clone or isolate | Reference | Inc type | Size (bp) | No. AMR genes | AMR genotype | Metal resistance |
| --- | --- | --- | --- | --- | --- | --- | --- |
| pST34VN2 | VN2 | This study | IncA/C2 | 173,870 | 11 | *floR, catA2, sul2, tetA, strAB, aac(3)-IId, qnrS1, bla<sub>CTX-M-55</sub>, bla<sub>TEM</sub>, mcr3.1* | *merADE* |
| pST34VN4 | VN4 | This study | IncA/C2 | 175,360 | 11 | *floR, sul2, tetA, strAB, aac(3)-IId, qnrS1, mcr3.1, ermF, ereD, mphA* |  |
| pST34VN3 | VN3 | This study | IncHI2 | 247,148 | 15 | *floR, cmlA, catA2, sul2, dfrA14, strAB, aadA, aph(3')-Ia, aac(3)-IId, qnrS1, bla<sub>TEM</sub>, mphA, lnu(F), arr-2* | *terWZBCD* |
| pVNB151 (V-BSI) | VBSI | Mather et al | IncHI2 | 246,444 | 16 | *floR, cmlA, catB3, sul1, sul2, sul3, aph(3')-Ia, aadA, aac(6')-Iaa, aph(4)-Ia, aac(3)-IVa, oqxAB, aac(6')-lb-cr, bla<sub>OXA-1</sub>, arr-3* | *terWZBCD* |
| p10907_1 | 01_0907 (BAPS-4) | This study | IncFIA | 203,427 | 8 | *floR, cmlA, sul2, aadA1, aadA2, qnrS1, bla<sub>TEM</sub>, mcr1.1* |  |
| p10907_2 | 01_0907 (BAPS-4) | This study | IncFIB/<br>IncFII | 90,994 | 4 | *aadA2, aac(3)-IId, qnrS1, mcr3.1, lnu(F)* |  |

Likewise, the two IncA/C2 plasmids, isolated from VN2 (pST34VN2) and VN4 (pST34VN4), were similar in size (~170 kbp) but markedly different in genetic composition, including some AMR determinants (Table 3; Figure S5). While pST34VN2 carried more beta-lactamases (*bla*_CTX-M-55_, *bla*_TEM_), macrolide resistance genes (*mphA, ereD, ermF*) were enriched in pST34VN4. Phylogeny based on the IncA/C2 backbone demonstrated separate clustering of these two plasmids (Figure S6). pST34VN2 is genetically indistinguishable from the MDR plasmid (pAUSMDU00004549) found in the previously defined Australia Lineage 1 (BAPS-1)^10^ (Figures S5, S6). Notably, an IncFII transfer region (*traM* – *finO;* >39 kbp) was integrated into pST34VN2 backbone, which was not observed in pST34VN4. These findings suggest that pST34VN2 is highly conjugative and have been acquired by ST34 in several independent occasions. However, unlike its IncHI2 counterpart, the IncA/C2 replicon was restricted to only ST34 and not found in other *Salmonella* serotypes in Vietnam. We experimentally confirmed that pST34VN2 could be conjugally transferred to *E. coli* (conjugation frequency of 1.103E-04) and *Shigella sonnei* (4.862E-05), but not to *Salmonella* Kentucky. Taken together, these data support that the acquisition of large MDR plasmids (of IncHI2 or IncA/C2) contributed to the successful expansion of ST34 in Vietnam. Additionally, other plasmids (IncFIA, IncFIB, IncFIC, IncFII, IncHI1, IncI, IncX, IncR) were also found in ST34 collected in Vietnam, but they were less frequently associated with carriage of *bla*_CTX-M-55_ or *mph(A)*, and were not linked to major clonal expansion events. We also identified one isolate in Vietnam (01_0907, BAPS-4) that co-carried two plasmids (IncFIA, IncFIB/IncFII) encoding two different *mcr* variants (*mcr*-1.1 and -3.1 respectively) (Table 3).

### Global propagation of *Salmonella enterica* ST34 clones carrying *bla*_CTX-M-55_

The mobilization of pST34VN2-borne *bla*_CTX-M-55_ in ST34 poses serious public health concerns, so we sought to further investigate the extent of international spread pertaining to this variant. We queried recently compiled ST34 genome databases (n=9,589) for the presence of *bla*_CTX-M-55_ ^7^, and identified 91 new positive genomes. *bla*_CTX-M-55_ was also the predominant ESBL in the database (n=91/196). Addition of these genomes to the ST34 global phylogeny showed that *bla*_CTX-M-55_ was largely restricted to four clones (Figure S7), including the aforementioned VN2 and Australia Lineage 1 (Australia_L1). The co-transfer of *bla*_CTX-M-55_ and IncA/C2 plasmid (pST34VN2 and its derivatives) was notable in both these two clones, together with SEA_minor clone. High-confidence phylogenetic reconstructions suggest that the progenitors of VN2 and Australia_L1 might have emerged from animals in Southeast Asia, prior to their propagation in humans (Figure 5AB), though this needs further confirmation in larger datasets. Intercontinental spread was evident in these two clones, particularly with several probable introductions of VN2 from Southeast Asia to Australia, USA, China and UK. The phylogeny also displays the occasional deletions of the IncFII transfer region and *mcr*-3.1 in IncA/C2 plasmids. On the other hand, we identified a separate ST34 clone (situated within BAPS-4) mostly composed of sequences from China, and these were not shown to harbour any major plasmids (Figure 5D). This agrees with recent report of chromosome-borne *bla*_CTX-M-55_ *S*. 4,[5],12,i:-isolated from China^16^, and showed that this variant could have circulated endemically in animals and disseminated to high-income settings.

**Figure 5.**
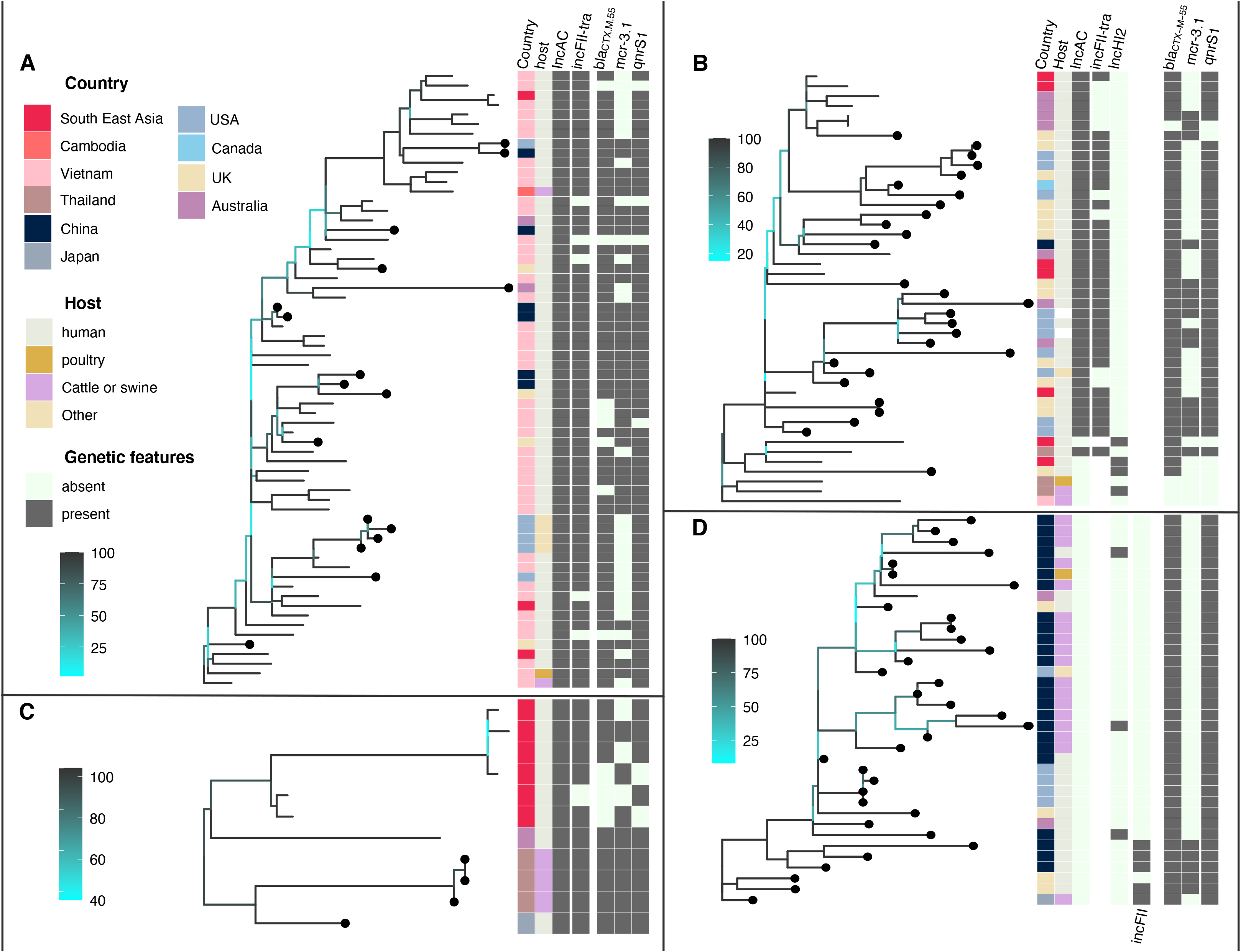
Phylogenetic structures of *Salmonella enterica* ST34 clones carrying *bla*_CTX-M-55_. Each panel displays a rooted phylogeny of a multidrug resistant clone, including (A) VN2, (B) Australia lineage 1 (Aus_L1), (C) Southeast Asia minor clone (SEA_min), and (D) China. Each phylogeny’s branches are coloured in accordance to bootstrapping values, from low (cyan) to high (black) (see legend). New included genomes, aside from those shown in Figure 1, are marked with filled black circles on tips. The appended heatmap displays data associated with each taxon, including original country of isolation, host of isolation, presence of plasmid replicons (IncA/C2, IncHI2, IncFII), IncFII-tra region incorporated on IncA/C2 plasmid, and resistance to critically important antimicrobials (*bla*_CTX-M-55_, *mcr-3.1, qnrS1*).

## Discussion

Our work offers a novel insight into the evolution and epidemiology of *S*. 4,[5],12,i:-ST34 in Southeast Asia. Using isolates sourced from surveillance studies, we revealed that ST34 has likely been introduced into Vietnam, most likely sourced from Europe, in at least nine occasions since the 2000s. This pattern and timing are similar to the multiple disseminations of ST34 from Europe to USA ^23^, highlighting the increased dynamic in global transmission of this pathogen at the start of the 21^st^ century. Our findings also underscore the role of Southeast Asia as a hub for intercontinental spread of *S*. 4,[5],12,i:-ST34, confirming a previous interpretation relying on isolates from Australian travellers to the region ^10^.

Importantly, the acquisition of large MDR plasmids, of IncHI2 or IncA/C2, underpinned successful clonal expansions of ST34 in Vietnam, with a trend of increasing resistance to CIAs (fluoroquinolone, ceftriaxone, colistin, and azithromycin). This echoes previous reports of ESBL or *mcr* positive *S*. 4,[5],12,i:-in Asia ^15–17,34^, suggesting that these variants are potentially widespread. It is most likely that ST34 picked up MDR plasmids from the vast pool of local Enterobacterales, as demonstrated previously for the local circulation of other enteric pathogens such as *S. sonnei* ^35,36^. This phenomenon may occur more frequently in countries like Vietnam, where the total estimated antimicrobial usage for humans and animals is nearly twice that of the European Union ^37^. Among examined sectors, the swine industry bears the highest consumption of antimicrobials, thus creating a favourable environment for MDR *Salmonella* variants to thrive.

Consistent with other reports, IncHI2 plasmids were identified as the predominant vehicle encoding resistance to CIAs in ST34, as well as in other *Salmonella* sequence types ^38,39^. Diversity in genetic makeup and co-transferring AMR determinants, as revealed in the IncHI2 plasmid tree, indicate its ancestral origin and complex circulation in *Salmonella*. In contrast, though IncA/C2 plasmids were previously recovered from several *Salmonella* serotypes ^40^, we found that it circulated exclusively in ST34 in Vietnam. The widespread propagation of IncA/C2 plasmids has raised public health concerns since they frequently carry the ESBL gene *bla*_CMY-2_, particularly in *S*. Typhimurium and *S*. 4,[5],12,i:-^23^.

This *bla*_CMY-2_ carrying IncA/C2 shares many genetic similarities with pST34VN2 described herein, including AMR (*floR, tetA, sul2, strAB*) and mercury resistance genes (*merADE*). However, pST34VN2 co-transfers numerous CIA resistance elements (*qnrS1, mcr-3.1*, and *bla*_CTX-M-55_), contributing to the regional and intercontinental expansions of several ST34 clones. Similar to other *mcr* variants, *mcr-3.1* most probably originated from animals ^41^, and *bla*_CTX-M-55_ was reported as the most prevalent ESBL originated from *E. coli* in animals in Vietnam ^42,43^. Together with evidence from phylogenetics, it is speculated that pST34VN2 likely first evolved in animals. In agreement with previous findings ^16^, we confirmed that the plasmid is self-conjugative to *E. coli* and *S. sonnei*, raising the concern of its wide propagation in local Enterobacterales pools. It was shown that high dose treatment of tetracycline nearly quadrupled the conjugation frequency of IncA/C2 plasmid in vivo ^44^, and this antibiotic ranks among the most frequently used in husbandry in Southeast Asia ^45,46^. This observation, together with IncFII conjugation machinery incorporated into pST34VN2, could greatly accelerate this plasmid’s expansion. Additionally, the finding that pST34VN2 failed to transfer to *S*. Kentucky was likely due to high fitness cost associated with retaining the IncA/C2 ^47^. *In vivo* experiments demonstrated that IncA/C2 incurred the lowest fitness cost in *S*. Typhimurium and its monophasic variant recipients, but was deleterious to growth of other *Salmonella* serotypes^44^. Our findings also support this notion, evidenced by the intercontinental expansion of two separate clones carrying pST34VN2 (VN2 and Australia_L1), as well as the plasmid’s sustained inheritance in these genetic backgrounds.

A novel contribution of this study is using mutation analysis to inspect the evolutionary trajectory and adaptation of ST34 at the global scale. The most notable genes identified under selection include ones related to antimicrobial response, virulence and host interactions. Polymyxins and quaternary ammonium compounds are frequently used in husbandry as prophylactic antibiotic and farm disinfectants ^48^, of which pressure could have favoured the positive selection observed in *arnA* and *smvA*, respectively. This, together with the recorded diversity of *mcr* variants found in ST34, points to its enhanced adaptation under exposure to colistin. Regarding virulence, ST34 exhibited a lesser extent of systemic dissemination due to the loss of *S*. Typhimurium virulence plasmid ^49^. Accumulation of mutations in several virulence factors (*phoQ, misL, barA, fliC, cspC*) suggest that ST34 has evolved to reinvent its interactions with host cells. Experimental evidences demonstrated *S*. 4,[5],12,i:-ST34 survived better in human macrophages compared to *S*. Typhimurium ^49^, and this effect was lineage-dependent, with higher intracellular bacterial replication rate observed in BAPS-3 and 4 ^10^. However, we reported here that the intra-macrophage virulence activator *cspC* was the most frequent target for degradation, implying that ST34 evolved toward dampened virulence and host invasion. Compared to *S*. Typhimurium U288, ST34 inoculation in pigs resulted in bacterial loads that were greater in faeces, but lower in mesenteric lymph nodes ^50^. Besides, the strong negative selection identified in *bapA* suggests that preserving biofilm formation is crucial for ST34 adaptation. Indeed, ST34 recovered from China were shown to form stronger biofilms compared concurrent *S*. Typhimurium ST19 ^51^. We compared the findings from ST34 with those deduced from fluoroquinolone resistant *S. sonnei*, an enteric pathogen sharing similar evolution timeframe and international spread ^52^. There is little overlap between the two species in the repertoire of genes under potential selection, except for the general trend in response to antimicrobials. ST34 also harboured fewer lineage-defining pseudogenes, reflecting its more versatile lifestyle compared to the obligate intracellular pathogen *S. sonnei*. Stepwise pseudogenizations have been characterized in details for the BSI-causing *S*. Typhimurium ST313, highlighting that reductive evolution was associated with more invasive disease manifestation ^4^. We examined the occurrence of ST313-defining pseudogenes in our ST34 collection, and found that none of these has been maintained in the evolution of ST34. Interestingly, the divergence of VBSI ST34 was preceded by fixation of a frameshift mutation in *pduT*, involved in utilization of 1,2-propanediol ^22^. Fermentation of the microbiota-derived 1,2-propanediol facilitates *Salmonella* expansion in the gastrointestinal tract ^53^, and disruptions in this metabolic pathway have been documented to serotypes causing more invasive illnesses ^54^. It is likely that *pduT* mutation was among the initial steps to facilitate extraintestinal adaptation. Despite its detection in cattle and poultry, VBSI was not attributed to pediatric diarrhea, indicating that this variant may be outcompeted for causing gastroenteritis or restricted to circulation in specific vulnerable populations.

Genomic surveillance of ST34 has been implemented frequently in developed settings, where sample collection and sequencing are centralized. The paucity of genomic data from resource-limited but epidemiologically significant regions, like Southeast and South Asia, poses major gaps in understanding and tracking the molecular evolution of this widespread genotype. Our study highlighted Southeast Asia as an important hotspot for emergence and transmission of MDR ST34 *S*. 4,[5],12,i:-, calling for strengthening surveillance efforts in this region for such important NTS genotypes. *Salmonella* resistant to CIAs, such as ceftriaxone or azithromycin, were associated with prolonged hospitalization in young children with gastroenteritis ^21^, and could lead to treatment failures in vulnerable patients ^55^. Close attention should also be paid to ST34 causing invasive diseases (VBSI clone), in light of the evolving HIV epidemic in Southeast Asia ^56^. Generally, our work greatly complements ongoing global efforts in elucidating the epidemiology and evolution of ST34, and generated evidence for future intervention strategies.

## Materials and Methods

### Organism collection and whole genome sequencing

This study focused on investigating the genomic epidemiology *of Salmonella enterica* ST34, of both serotypes Typhimurium and 4,[5],12,i:-(monophasic variant), in Vietnam and Southeast Asia. For Vietnamese sequences, we combined 77 ST34 genomes published previously ^22^ with 133 isolates sequenced for this study. We selected representative isolates from a recent genomic study of ST34 in Australia (179/309) for global phylogenetic context, covering different lineages, geographies and isolation times ^10^. Additionally, we included sequences from other Asian countries (maximum 15 genomes for each), accessed via the Enterobase database ^57^. The resulting compiled data include 454 *S. enterica* ST34 sequences (Vietnam, n=210; Cambodia, n=4; Thailand, n=15; Laos, n=4; Southeast Asia, n=40; mainland China, n=14; Japan, n=13; Taiwan, n=15; Australia, n=38; UK, n=56; Italy, n=13; USA, n=32). Isolates from Vietnam were mostly sourced from a paediatric diarrheal surveillance study^21^ and a bloodstream infection investigation in Southern Vietnam (for isolates of human origin), as well as from faecal material of asymptomatic animals (poultry and swine). For isolates recovered from patients with recent travel history (Australian dataset), the travel destination was recorded as the geography of origin for these isolates. For 133 Vietnamese isolates sequenced for this study, genomic DNA was extracted from pure culture colonies using the Wizard genomic DNA extraction kit (Promega, USA) and sent to the Wellcome Sanger Institute for whole genome sequencing using the Illumina Hiseq 2500 platform, generating paired-end reads (125bp ×2).

### Short read mapping and phylogenetic reconstruction

For all isolates, sequencing quality was assessed using FastQC v0.11.5, showing that the compiled dataset comprised sequence reads of varying lengths (75 – 300 bp). Trimmomatic v0.38 was used to remove adapters and filter reads with low sequencing quality (SLIDINGWINDOW:10:22, HEADCROP:10 – 15), dependent on the corresponding read length of each isolate (MINLEN: 35 – 50). All trimmed read pairs were mapped against the recently published complete genome of ST34 serotype 4,[5],12,i:-(TW-Stm6, accession number: CP019649) using BWA-mem v0.7.17 with default settings, and duplicate reads were removed using PICARD, followed by indel realignment using GATK v3.7.0 ^58^. We further removed reads with nonoptimal local alignment using samclip (https://github.com/tseemann/samclip) to avoid false positives during variant calling. Single nucleotide variants (SNVs) were identified using the haplotype-based caller Freebayes v1.3.6, and low quality SNPs were removed using bcftools v1.12 if they met any of these criteria: consensus quality < 30, mapping quality < 30, read depth < 4, ratio of SNVs to reads at a position (AO/DP) < 85%, coverage on the forward or reverse strand < 1 ^59,60^. For each isolate, a pseudogenome (same length as reference) was created using the bcftools ‘consensus’ command, incorporating the filtered SNVs and invariant sites while masking regions with low mapping (depth < 4) and low-quality SNVs with ‘N’. An alignment of 454 *S. enterica* ST34 isolates was generated, and we further masked regions pertaining to insertion sequences, transposases, prophages (predicted by PHASTER), and recombination (as detected by Gubbins v1.4.5) ^61,62^. Subsequently, invariant sites were removed, producing an SNP alignment of 4,962 bp. This was input into RAxML v8.2.4 to reconstruct a maximum likelihood phylogeny under the GTRGAMMA model with 500 bootstrap replicates ^63^. This phylogeny was rooted using a Vietnamese *S. enterica* ST19 isolate. The R package ggtree v3.2.1 was used to append associated metadata to the phylogeny ^64^. Additionally, the SNP alignment was input into Fastbaps v1.0.6 for hierarchical Bayesian analysis of population structure in a phylogeny-independent approach ^65^.

### Bayesian phylogenetic inference

In order to analyse the temporal structure of *S. enterica* ST34, we subsampled our collection to include 222 representative isolates, based on the location and time of isolation, resulting in an SNP alignment of 2,671 bp. Phylogenetic reconstruction was repeated for these isolates, following aforementioned procedures. This generated a phylogeny with similar topology and representativeness to that of the original full dataset (n=454). TempEst v1.5.1 was utilized to estimate the linear relationship between the resulting phylogeny’s root-to-tip divergence and the isolates’ sampling dates (in month/year)^66^, which indicated the presence of modest temporal structure (R^2^=0.33). Bayesian phylogenetic inferences were conducted using BEAST v1.10.4 to estimate *S. enteric* ST34’s substitution rate and tMRCAs of clonal expansions in Vietnam^67^. In order to identify the best suited model for this dataset, we conducted triplicate BEAST runs on a number of combinatory models. These include ones with a GTR + Γ4 substitution model, with either a strict or relaxed lognormal clock model, in conjunction with a constant or exponential growth demographic model. Each analysis was performed using a continuous 100 – 150 million generation MCMC chain, with samples taken every 10,000 – 15,000 generations, respectively. Parameter convergences were visually assessed using Tracer v1.7 (ESS > 200 for all parameters). For each analysis, both path sampling and stepping-stone sampling approaches were implemented to estimate the marginal likelihood ^68^. The best model, as selected based on the comparison of calculated Bayes factors, was a relaxed lognormal clock model with an exponential growth demographic model. We combined this model’s triplicate runs using LogCombiner and TreeAnnotator v1.10.4, with 20% burnin removal, and output the maximum clade credibility (MCC) tree and inferred parameters.

### Phylogeography analysis of ST34

We combined the maximum likelihood phylogeny of 454 ST34 genomes with the isolates’ geographical location (subcontinental level) to infer the degree of intercontinental spread of this pathogen. To reduce bias inherent to unequal sampling, we subsampled the original phylogeny to include equal numbers of isolates from each geography (n=30 each from America, East Asia, Europe, Oceania, and Southeast Asia), generating 1,000 subsampled trees. Stochastic mapping, as implemented in the function *make.simmap* in the R package phytools v1.0-1, was performed on each subsampled tree to quantify the number of transition events between geographical states and the proportional evolutionary time spent within each geography ^69,70^. The analysis was conducted under an asymmetric model of character change (ARD) with the rate matrix sampled from the posterior probability distribution using MCMC (Q=mcmc) for 100 simulations. In addition, we permuted (without replacement) the location of the original phylogeny to create ten randomization sets, which were independently subjected to stochastic mapping (as aforementioned with 500 subsampled trees). Analysis of variance (ANOVA) with post-hoc Tukey test were used to compare the results from ‘true’ and ‘randomization’ runs.

### Determination of the accessory genome

For each isolate, trimmed paired-end reads were input into Unicycler v0.4.9 to generate a de novo assembly ^71^. Annotation was determined by Prokka v1.14.6 for each assembly, using the TW-Stm6 Genbank file as the reference ^72^. The presence of acquired AMR genes and plasmid incompatibility types on each assembly were detected by running Abricate v0.7, with references to the curated ResFinder and PlasmidFinder databases, respectively ^73,74^. ABACAS v1.3.1 was used to order the genome assembly of ST34 to the TW-Stm6 reference, and non-aligned contigs (predicted to belong to plasmids) were queried against the public database using BLASTN or the web version of PLSDB ^75,76^.

### dN/dS analysis on substitutions

Substitution mutations, as identified by mapping to the TW-Stm6 reference, were summarized and annotated using SnpEff v5.0e ^77^. Mutations present within mobile, repetitive, or recombination regions (as defined in the ‘Phylogenetic reconstruction’ section) were filtered out, resulting in a collection of 4,962 SNPs for investigation. We adopted a previously published approach to assess genewise dN/dS ratio (non-synonymous to synonymous substitution rate) ^78^. Briefly, ancestral state reconstruction for all SNPs were conducted using PAML, based on the input alignment and maximum likelihood phylogeny ^79^. Mutations were classified as intergenic, synonymous, and non-synonymous by comparing each SNP’s annotation to the reconstructed ancestral state. The dN/dS ratio was adjusted for transition/tranversion rate and codon usage under the NY98 model. Genes were determined as undergoing positive selection if their adjusted dN/dS ratio was > 2 or if they had no synonymous and at least five non-synonymous mutations.

### Plasmid sequencing and plasmid core phylogeny

In order to determine the full-length sequence of multidrug resistance plasmids associated with ST34’s successful clonal expansions in Vietnam, we selected three representative isolates (01_0119: VN2; 02_1644: VN3; 02_1206: VN4) for long-read plasmid sequencing. Plasmid DNA was extracted from overnight culture using the Plasmid Midi kit (QIAGEN, Germany), following manufacturer’s instructions. The purified DNA was input into the Rapid Barcoding kit (Oxford Nanopore Technology, SQK-RBK004) for sample multiplexing and library preparation, and the resulting libraries were sequenced on a Flongle flow cell R9.4.1 (Oxford Nanopore Technology, United Kingdom). Base calling and demultiplexing were carried out using Guppy v6.3.8 (https://community.nanoporetech.com), and hybrid assembly was conducted for each isolate using Unicycler v0.4.9 on the filtered long-read and corresponding Illumina short-read sequences. The reconstructed plasmid sequences were visualized and compared using Easyfig^80^.

We further screened the presence of MDR plasmids of IncA/C and IncHI2 types on the genome databases of *S. enterica* of other genotypes, recovered from the same diarrheal surveillance study. Sequencing reads of isolates carrying IncA/C (n=89) and IncHI2 (n=143) plasmids were mapped to the full-length references AUSMDU00004549 plasmid P01 (NZ_OU015324.1; isolated in Australia) and p16-6773 (NZ_CP039861; isolated in Canada), respectively. The former reference is homologous to pST34VN2 except for the deletion of the IncFII transfer region. Regions pertaining to IS elements and recombination, as detected from Gubbins v1.4.5 (10 iterations), as well as invariant sites were removed, producing alignments of 62 (for IncA/C) and 328 SNPs (for IncHI2). Maximum likelihood phylogenies were reconstructed from these alignments using RAxML v8.2.4, with 500 bootstraps.

### Conjugation experiments

Conjugation was performed between the ciprofloxacin susceptible ESBL-positive *S*. 4,[5],12,i:-strain 01_0119 (VN2 clone, donor) and the ciprofloxacin resistant ESBL-negative *E. coli* CTH01, *Shigella sonnei* 03_0520, and *Salmonella* Kentucky ST198 01_0211 (clinical isolates, recipients). For each bacterial strain, 50μL overnight culture was incubated in 5 ml Luria-Bertani (LB) broth (supplemented with appropriate antibiotics) and grown at 37 °C to an optical density (OD_600nm_) of 0.3. Then, 50μL of each donor and recipient cultures were combined in 5 ml of LB broth and grown without shaking for 24 hours at 37 °C. Transconjugants were selected on MacConkey plates supplemented with ciprofloxacin (4 mg/l) and ceftriaxone (6 mg/l), while donors were selected and enumerated on MacConkey plates supplemented with ceftriaxone (6 mg/l). The experiment was conducted three times for each donor-recipient combination, and conjugation frequency was calculated as the number of transconjugants per donor cells.

### Investigation on ST34 carrying *bla*_CTX-M-55_

Our results indicated that carriage of *bla*CTX-M-55 was associated with a clonal expansion in Southeast Asian countries. We next sought to investigate the international spread and molecular epidemiology of ST34 bearing *bla*_CTX-M-55_ by querying a recently compiled genome database of *S. enterica* ST34 (n=9,589) ^7^. *Bla*_CTX-M-55_ was detected in 91 genomes, which were downloaded from the NCBI public database. Since *bla*_CTX-M-55_ is often co-transferred with *mcr-3.1*, we additionally included another sixteen *mcr-3.1* positive genomes, compiled from a recently published study in China ^16^. We combined these 107 public genomes with 222 representative assemblies in our study (used for temporal phylogenetics), and performed pangenome analysis using Panaroo (strict mode) ^81^. The resulting core genome (4,393 genes present in at least 95% of genomes) was aligned, and recombination was detected and removed using Gubbins v1.4.5 (ten iterations), producing an SNP alignment of 3,023 bp. This served as the input for phylogenetic reconstruction using RAxML v8.2.4, as described above. To further refine the phylogeny of individual clones of interest, we implemented reference-based mapping approach to isolates belonging to clones VN2, Australia_L1, SEA_minor, and China_ESBL. In cases where raw sequencing reads were not available from the NCBI database, simulated paired-end reads were generated from the isolate’s assembly using fastaq v3.17.0 (https://github.com/sanger-pathogens/Fastaq; to_perfect_reads: mean insert size=400, insert standard deviation=25, mean coverage=80, read length=125). Reads were mapped to the reference TW-Stm6, and SNP calling and phylogenetic reconstruction followed procedures described above.

## Data availability

Raw sequence data are available in the European Nucleotide Archive (ENA) under the project number PRJEB9121. Table S1 provides the detailed accession numbers and metadata for all genomes used in this study.

## Acknowledgements

HCT is a Wellcome International Training Fellow (218726/Z/19/Z). AEM is supported by the Biotechnology and Biological Sciences Research Council (BBSRC) through the BBSRC Institute Strategic Programme Microbes in the Food Chain BB/R012504/1 and its constituent project BBS/E/F/000PR10348 (Theme 1, Epidemiology and Evolution of Pathogens in the Food Chain). SB is a Wellcome Senior Research Fellow (215515/Z/19/Z). DTP is supported by the Wellcome International Training Fellowship (222983/Z/21/Z).

The authors wish to thank all study participants and their guardians for participating in the study.

## Referances

1. Kirk, M. D. et al. World Health Organization Estimates of the Global and Regional Disease Burden of 22 Foodborne Bacterial, Protozoal, and Viral Diseases, 2010: A Data Synthesis. PLoS Med. 12, 1–21 (2015).

2. Stanaway, J. D. et al. The global burden of non-typhoidal salmonella invasive disease: a systematic analysis for the Global Burden of Disease Study 2017. Lancet Infect. Dis. 19, 1312–1324 (2019).

3. Bawn, M. et al. Evolution of salmonella enterica serotype typhimurium driven by anthropogenic selection and niche adaptation. PLoS Genet. 16, 1–29 (2020).

4. Pulford, C. V et al. Stepwise evolution of Salmonella Typhimurium ST313 causing bloodstream infection in Africa. Nat. Microbiol. 6, 327–338 (2021).

5. Hopkins, K. L. et al. Multiresistant Salmonella enterica serovar 4,[5],12:i:- in Europe: A new pandemic strain? Eurosurveillance 15, 1–9 (2010).

6. Wong, M. H. Y. et al. Expansion of salmonella enterica serovar typhimurium ST34 clone carrying multiple resistance determinants in China. Antimicrob. Agents Chemother. 57, 4599–4601 (2013).

7. Trachsel, J. M., Bearson, B. L., Brunelle, B. W. & Bearson, S. M. D. Relationship and distribution of Salmonella enterica serovar I 4,[5],12:i:- strain sequences in the NCBI Pathogen Detection database. BMC Genomics 23, 1–21 (2022).

8. European Centre for Disease Prevention and Control European Food Safety Authority. Multi□country outbreak of monophasic Salmonella Typhimurium sequence type (ST) 34 linked to chocolate products – 12 April 2022. EFSA Supporting Publications 19, (2022).

9. Arai, N. et al. Identification of a Recently Dominant Sublineage in Salmonella 4,[5],12:i:- Sequence Type 34 Isolated From Food Animals in Japan. Front. Microbiol. 12, 1–12 (2021).

10. Ingle, D. J. et al. Evolutionary dynamics of multidrug resistant Salmonella enterica serovar 4,[5],12:i:- in Australia. Nat. Commun. 12, 1–13 (2021).

11. Elnekave, E. et al. Salmonella enterica Serotype 4,[5],12:i:-in Swine in the United States Midwest:An Emerging Multidrug-Resistant Clade. Clin. Infect. Dis. 66, 877–855 (2018).

12. Petrovska, L. et al. Microevolution of monophasic Salmonella typhimurium during epidemic, United Kingdom, 2005-2010. Emerg. Infect. Dis. 22, 617–624 (2016).

13. Mu, Y. et al. Genomic Epidemiology of ST34 Monophasic Salmonella enterica Serovar Typhimurium from Clinical Patients from 2008 to 2017 in Henan, China. Engineering (2022). doi:10.1016/j.eng.2022.05.006

14. Branchu, P. et al. SGI-4 in monophasic Salmonella typhimurium ST34 is a novel ice that enhances resistance to copper. Front. Microbiol. 10, 1–12 (2019).

15. Nadimpalli, M. et al. CTX-M-55-type ESBL-producing Salmonella enterica are emerging among retail meats in Phnom Penh, Cambodia. J. Antimicrob. Chemother. 74, 342–348 (2019).

16. Sun, R. Y. et al. Global clonal spread of mcr-3-carrying MDR ST34 Salmonella enterica serotype Typhimurium and monophasic 1,4,[5],12:i:- variants from clinical isolates. J. Antimicrob. Chemother. 75, 1756–1765 (2020).

17. Monte, D. F. et al. Multidrug-and colistin-resistant salmonella enterica 4,[5],12:i:-sequence type 34 carrying the mcr-3.1 gene on the IncHI2 plasmid recovered from a human. J. Med. Microbiol. 68, 986–990 (2019).

18. World Health Organization. WHO list of Critically Important Antimicrobials (CIA) 6th Revision. (2019).

19. Patra, S. D., Mohakud, N. K., Panda, R. K., Sahu, B. R. & Suar, M. Prevalence and multidrug resistance in Salmonella enterica Typhimurium: an overview in South East Asia. World J. Microbiol. Biotechnol. 37, 1–17 (2021).

20. Duong, V. T. et al. No Clinical benefit of empirical antimicrobial therapy for pediatric diarrhea in a high-usage, high-resistance setting. Clin. Infect. Dis. 66, 504–511 (2018).

21. Duong, V. T. et al. Genomic serotyping, clinical manifestations, and antimicrobial resistance of non-typhoidal Salmonella gastroenteritis in hospitalized children in Ho Chi Minh City, Vietnam. J. Clin. Microbiol. 58, 1–15 (2020).

22. Mather, A. E. et al. New variant of multidrug-resistant Salmonella enterica serovar typhimurium associated with invasive disease in immunocompromised patients in Vietnam. MBio 9, 1–11 (2018).

23. Elnekave, E. et al. Transmission of multidrug-resistant salmonella enterica subspecies enterica 4,[5],12:i:-Sequence type 34 between europe and the United States. Emerg. Infect. Dis. 26, 3034–3038 (2020).

24. Baylay, A. J., Piddock, L. J. V. & Webber, M. A. Molecular mechanisms of antibiotic resistance revisited. Nat. Rev. Microbiol. 1–26 (2022). doi:10.1002/9781119593522.ch1

25. Gatzeva-Topalova, P. Z., May, A. P. & Sousa, M. C. Structure and mechanism of ArnA: Conformational change implies ordered dehydrogenase mechanism in key enzyme for polymyxin resistance. Structure 13, 929–942 (2005).

26. Carvalho, F. a, Aitken, J. D., Gewirtz, a T. & Vijay-Kumar, M. TLR5 activation induces secretory interleukin-1 receptor antagonist (sIL-1Ra) and reduces inflammasome-associated tissue damage. Mucosal Immunol. 4, 102–111 (2011).

27. Groisman, E. A. The pleiotropic two-component regulatory system PhoP-PhoQ. J. Bacteriol. 183, 1835–1842 (2001).

28. Altier, C., Suyemoto, M., Ruiz, A. I., Burnham, K. D. & Maurer, R. Characterization of two novel regulatory genes affecting Salmonella invasion gene expression. Mol. Microbiol. 35, 635–646 (2000).

29. Wang, S. et al. Autotransporter MisL of Salmonella enterica serotype Typhimurium facilitates bacterial aggregation and biofilm formation. FEMS Microbiol. Lett. 365, 1–8 (2018).

30. Latasa, C. et al. BapA, a large secreted protein required for biofilm formation and host colonization of Salmonella enterica serovar Enteritidis. Mol. Microbiol. 58, 1322–1339 (2005).

31. Bleibtreu, A. et al. The rpoS gene is predominantly inactivated during laboratory storage and undergoes source-sink evolution in Escherichia coli species. J. Bacteriol. 196, 4276–4284 (2014).

32. Cascales, E. et al. Colicin biology. Microbiol. Mol. Biol. Rev. 71, 158–229 (2007).

33. Choi, J., Salvail, H. & Groisman, E. A. RNA chaperone activates Salmonella virulence program during infection. Nucleic Acids Res. 49, 11614–11628 (2021).

34. Luo, Q. et al. MDR Salmonella enterica serovar Typhimurium ST34 carrying mcr-1 isolated from cases of bloodstream and intestinal infection in children in China. J. Antimicrob. Chemother. 75, 92–95 (2020).

35. Holt, K. E. et al. Tracking the establishment of local endemic populations of an emergent enteric pathogen. Proc. Natl. Acad. Sci. U. S. A. 110, 17522–7 (2013).

36. Pham, T. D. et al. Commensal Escherichia coli are a reservoir for the transfer of XDR plasmids into epidemic fluoroquinolone-resistant Shigella sonnei. Nat. Microbiol. 5, 1–9 (2020).

37. Carrique-mas, J. J., Choisy, M., Cuong, N. Van, Thwaites, G. & Baker, S. An estimation of total antimicrobial usage in humans and animals in Vietnam. Antimicrob. Resist. Infect. Control 7, 1–6 (2020).

38. Chen, W. et al. IncHI2 plasmids are predominant in antibiotic-resistant Salmonella isolates. Front. Microbiol. 7, 1–10 (2016).

39. Bloomfield, S. et al. Mobility of antimicrobial resistance across serovars and disease presentations in non-typhoidal Salmonella from animals and humans in Vietnam. Microb. Genomics 8, 1–13 (2022).

40. Hoffmann, M. et al. Comparative sequence analysis of multidrug-resistant incA/C plasmids from salmonella enterica. Front. Microbiol. 8, 1–13 (2017).

41. Yin, W. et al. Novel plasmid-mediated colistin resistance gene mcr-3 in Escherichia coli. MBio 8, 4–9 (2017).

42. Trung, N. V. et al. Limited contribution of non-intensive chicken farming to ESBL-producing Escherichia coli colonization in humans in Vietnam: An epidemiological and genomic analysis. J. Antimicrob. Chemother. 74, 561–570 (2019).

43. Nguyen, M. N. et al. Prospective One Health genetic surveillance in Vietnam identifies distinct blaCTX-M-harbouring Escherichia coli in food-chain and human-derived samples. Clin. Microbiol. Infect. 27, 1515.e1–1515.e8. (2021).

44. Johnson, T. J. et al. In Vivo transmission of an incA/C plasmid in Escherichia coli depends on tetracycline concentration, and acquisition of the plasmid results in a variable cost of fitness. Appl. Environ. Microbiol. 81, 3561–3570 (2015).

45. Van Cuong, N. et al. High-resolution monitoring of antimicrobial consumption in Vietnamese small-scale chicken farms highlights discrepancies between study metrics. Front. Vet. Sci. 6, 1–13 (2019).

46. Coyne, L. et al. Characterizing antimicrobial use in the livestock sector in three south east asian countries (Indonesia, thailand, and vietnam). Antibiotics 8, (2019).

47. Subbiah, M., Top, E. M., Shah, D. H. & Call, D. R. Selection pressure required for long-term persistence of blaCMY-2-positive IncA/C plasmids. Appl. Environ. Microbiol. 77, 4486–4493 (2011).

48. Kempf, I., Jouy, E. & Chauvin, C. Colistin use and colistin resistance in bacteria from animals. Int. J. Antimicrob. Agents 48, 598–606 (2016).

49. Sarichai, P. et al. Pathogenicity of clinical Salmonella enterica serovar Typhimurium isolates from Thailand in a mouse colitis model. Microbiol. Immunol. 64, 679–693 (2020).

50. Kirkwood, M. et al. Ecological niche adaptation of Salmonella Typhimurium U288 is associated with altered pathogenicity and reduced zoonotic potential. Commun. Biol. 4, 1–15 (2021).

51. Li, W.et al. Clonal expansion of biofilm-forming Salmonella Typhimurium ST34 with multidrug-resistance phenotype in the Southern Coastal Region of China. Front. Microbiol. 8, 1–6 (2017).

52. Chung The, H. et al. Dissecting the molecular evolution of fluoroquinolone-resistant Shigella sonnei. Nat. Commun. 10, 48–28 (2019).

53. Faber, F. et al. Respiration of Microbiota-Derived 1,2-propanediol Drives Salmonella Expansion during Colitis. PLoS Pathog. 13, 1–19 (2017).

54. Nuccio, S. P. & Bäumler, A. J. Comparative analysis of Salmonella genomes identifies a metabolic network for escalating growth in the inflamed gut. MBio 5, (2014).

55. Thanh, D. P. et al. A novel ciprofloxacin-resistant subclade of H58 Salmonella Typhi is associated with fluoroquinolone treatment failure in Nepal. Elife Under Review. (2016). doi:10.7554/eLife.14003

56. Van Griensven, F. et al. The continuing HIV epidemic among men who have sex with men and transgender women in the ASEAN region: Implications for HIV policy and serVICe programming. Sex. Health 18, 21–30 (2021).

57. Achtman, M. et al. Genomic diversity of Salmonella enterica-The UoWUCC 10K genomes project. Wellcome Open Res. 5, 1–22 (2021).

58. McKenna, A. et al. The Genome Analysis Toolkit: A MapReduce framework for analyzing next-generation DNA sequencing data. Genome Res. 20, 1297–303 (2010).

59. Garrison, E. & Marth, G. Haplotype-based variant detection from short-read sequencing. Preprint at https://doi.org/10.48550/arXiv.1207.39 (2012).

60. Danecek, P. et al. Twelve years of SAMtools and BCFtools. Gigascience 10, 1–4 (2021).

61. Arndt, D. et al. PHASTER: a better, faster version of the PHAST phage search tool. Nucleic Acids Res. 1–6 (2016). doi:10.1093/nar/gkw387

62. Croucher, N. J. et al. Rapid phylogenetic analysis of large samples of recombinant bacterial whole genome sequences using Gubbins. Nucleic Acids Res. 44, 1–13 (2014).

63. Stamatakis, A. RAxML version 8: a tool for phylogenetic analysis and post-analysis of large phylogenies. Bioinformatics 30, 1312–3 (2014).

64. Yu, G., Smith, D. K., Zhu, H., Guan, Y. & Lam, T. T. Y. ggtree: An r package for visualization and annotation of phylogenetic trees with their covariates and other associated data. Methods Ecol. Evol. 8, 28–36 (2016).

65. Tonkin-Hill, G., Lees, J. A., Bentley, S. D., Frost, S. D. W. & Corander, J. Fast hierarchical Bayesian analysis of population structure. Nucleic Acids Res. 47, 5539–5549 (2019).

66. Rambaut, A., Lam, T. T., Max Carvalho, L. & Pybus, O. G. Exploring the temporal structure of heterochronous sequences using TempEst (formerly Path-O-Gen). Virus Evol. 2, vew007 (2016).

67. Drummond, A. J. & Rambaut, A. BEAST: Bayesian evolutionary analysis by sampling trees. BMC Evol. Biol. 7, 214 (2007).

68. Baele, G., Li, W. L. S., Drummond, A. J., Suchard, M. A. & Lemey, P. Accurate model selection of relaxed molecular clocks in Bayesian phylogenetics. Mol. Biol. Evol. 30, 239–243 (2013).

69. Huelsenbeck, J. P., Nielsen, R., Bollback, J. P., Biology, S. & Apr, N. Stochastic Mapping of Morphological Characters Stochastic Mapping of Morphological Characters. Syst. Biol. 52, 131–158 (2003).

70. Revell, L. J. phytools: an R package for phylogenetic comparative biology (and other things). Methods Ecol. Evol. 3, 217–223 (2012).

71. Wick, R. R., Judd, L. M., Gorrie, C. L. & Holt, K. E. Unicycler: Resolving bacterial genome assemblies from short and long sequencing reads. PLoS Comput. Biol. 13, 1–22 (2017).

72. Seemann, T. Prokka: rapid prokaryotic genome annotation. Bioinformatics 30, 2068–9 (2014).

73. Zankari, E. et al. Identification of acquired antimicrobial resistance genes. J. Antimicrob. Chemother. 67, 2640–4 (2012).

74. Carattoli, A. et al. In Silico detection and typing of plasmids using plasmidfinder and plasmid multilocus sequence typing. Antimicrob. Agents Chemother. 58, 3895–3903 (2014).

75. Galata, V., Fehlmann, T., Backes, C. & Keller, A. PLSDB: a resource of complete bacterial plasmids. Nucleic Acids Res. 47, D195–D202 (2019).

76. Assefa, S., Keane, T. M., Otto, T. D., Newbold, C. & Berriman, M. ABACAS: Algorithm-based automatic contiguation of assembled sequences. Bioinformatics 25, 1968–1969 (2009).

77. Cingolani, P. et al. A program for annotating and predicting the effects of single nucleotide polymorphisms, SnpEff. Fly (Austin). 6, 80–92 (2012).

78. Lieberman, T. D. et al. Parallel bacterial evolution within multiple patients identifies candidate pathogenicity genes. Nat. Genet. 43, 1275–1280 (2011).

79. Yang, Z. PAML 4: Phylogenetic analysis by maximum likelihood.Mol. Biol. Evol. 24, 1586–1591 (2007).

80. Sullivan, M. J., Petty, N. K. & Beatson, S. A. Easyfig: A genome comparison visualizer. Bioinformatics 27, 1009–1010 (2011).

81. Tonkin-Hill, G. et al. Producing polished prokaryotic pangenomes with the Panaroo pipeline. Genome Biol. 21, 1–21 (2020).

