## Supplementary Information for "Multidrug resistance plasmids underlie clonal expansions and international spread of *Salmonella enterica* serotype 4,[5],12,i:- ST34 in Southeast Asia"

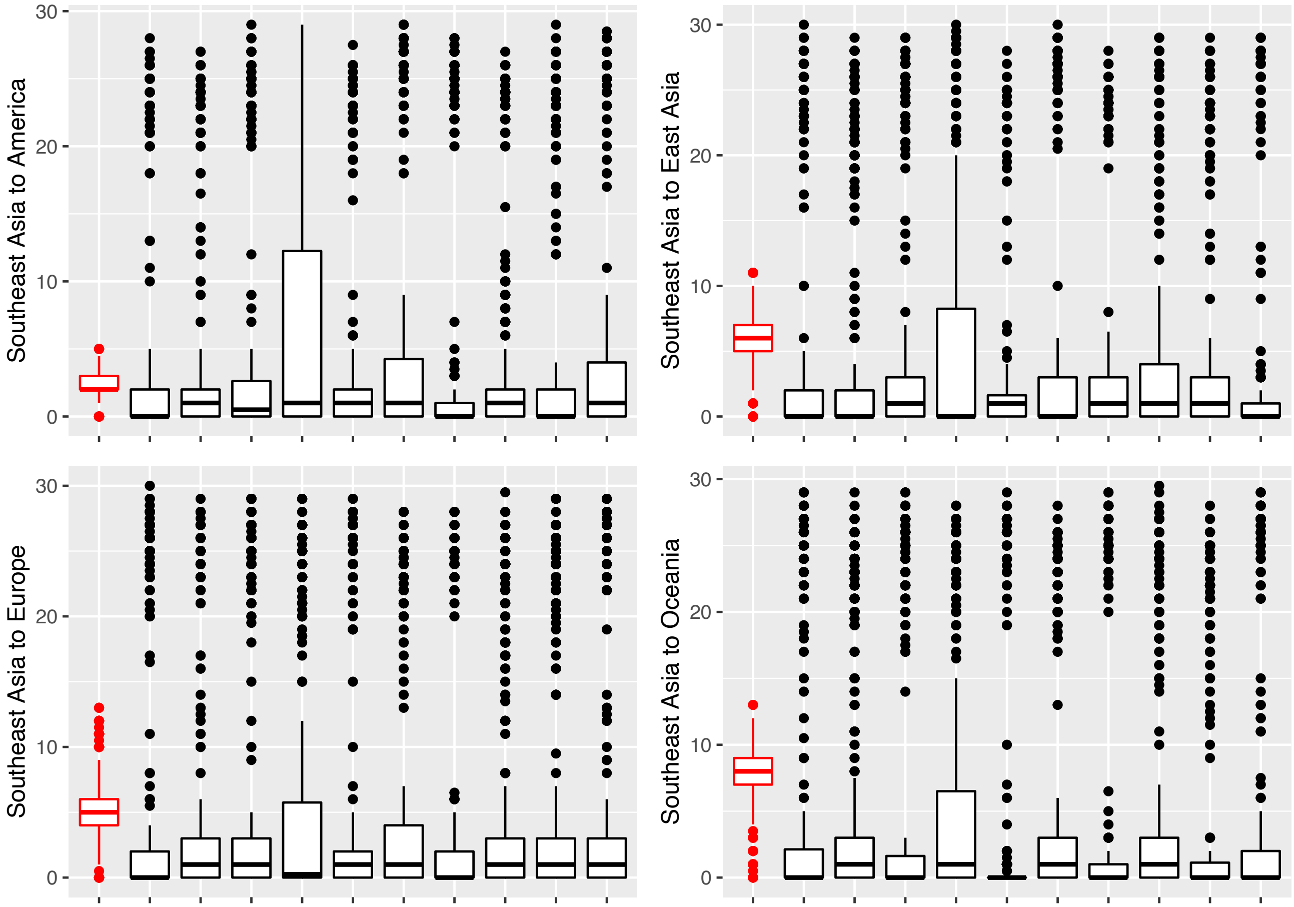


**Figure S1** Validation of phylogeographical signal in *Salmonella enterica* ST34. Stochastic mapping was performed on the original dataset (Figure 3) and ten tip-location randomized datasets (See Materials and Methods). Each panel summarizes the inferences made from the 'true' dataset (coloured red) and ten randomizations (coloured black), estimating the number of transition events from Southeast Asia to America, East Asia, Europe and Oceania during *Salmonella* ST34's evolutionary history. Each boxplot summarizes results from 1,000 (true dataset) and 500 (randomization) sub-samplings of the phylogenies.

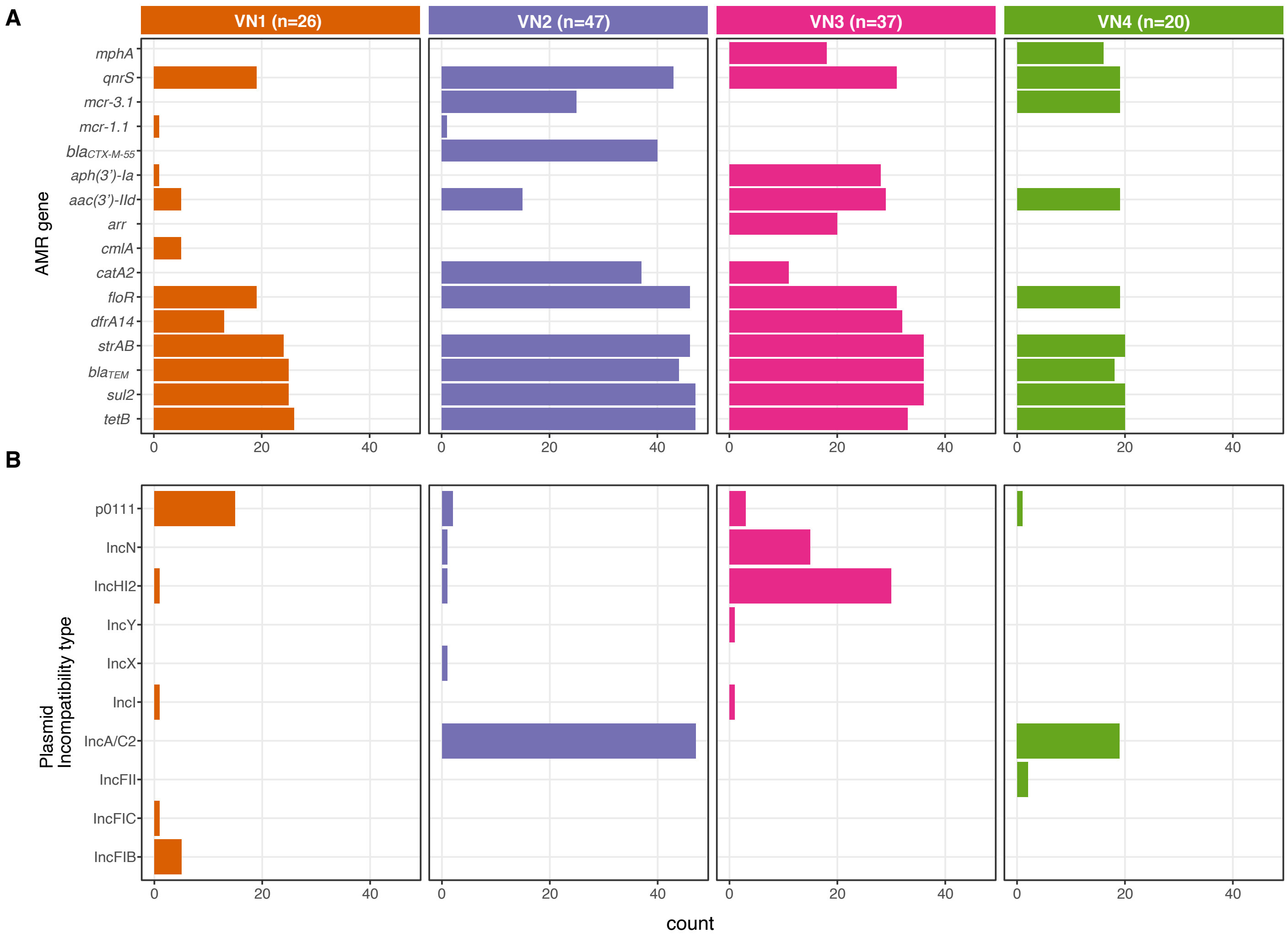


**Figure S2** Distribution of antimicrobial resistance (AMR) genes and plasmid types among four major *Salmonella enterica* ST34 clones causing gastroenteritis in Vietnam. For each panel, the bar graph displays the count of isolates from each clone (VN1 – VN4) carrying a respective element, stratified by (A) AMR genes, and (B) plasmid incompatibility types.

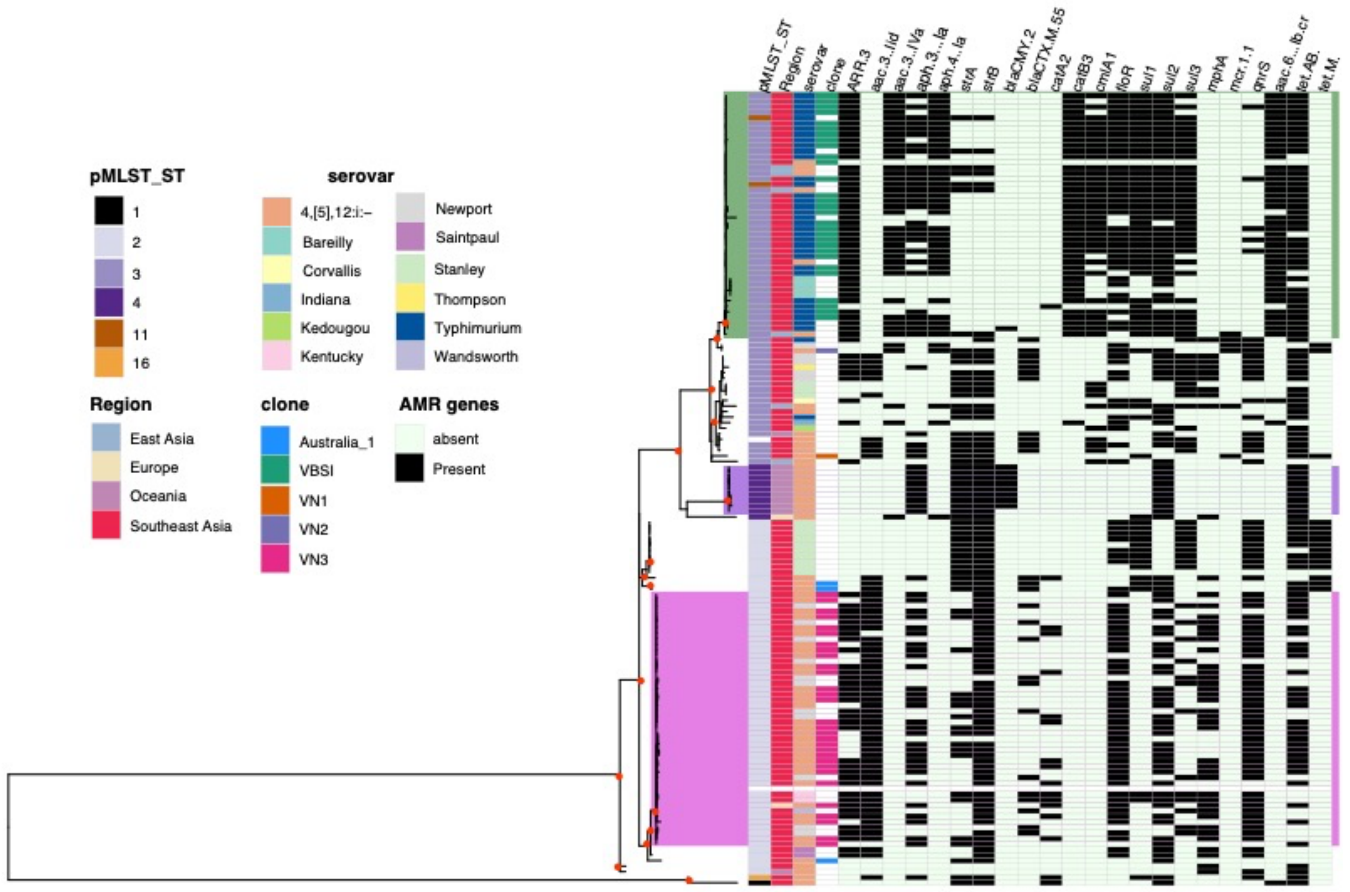


**Figure S3** Plasmid phylogeny of IncHI2 recovered from *Salmonella enterica*. The figure displays the maximum likelihood phylogeny of IncHI2 plasmids recovered from 143 *Salmonella enterica* isolates of diverse serotypes, constructed from 328 single nucleotide polymorphisms. The phylogeny is midpoint rooted. Filled red circles signify bootstrap support of ≥80 at internal nodes. The appended heatmap shows data associated with each taxon, including pMLST typing, region of isolation, predicted serotype based on genome sequencing, *Salmonella* ST34 clones, and presence of AMR genes. Coloured shades highlight plasmid clusters detected mainly in ST34, including green (VBSI, ST3, homologous to pVNB151), light magenta (VN3, ST2, homologous to pST34VN3), and dark magenta (ST4, endemic circulation in Australia, carrying *bla*$_{CMY-2}$).

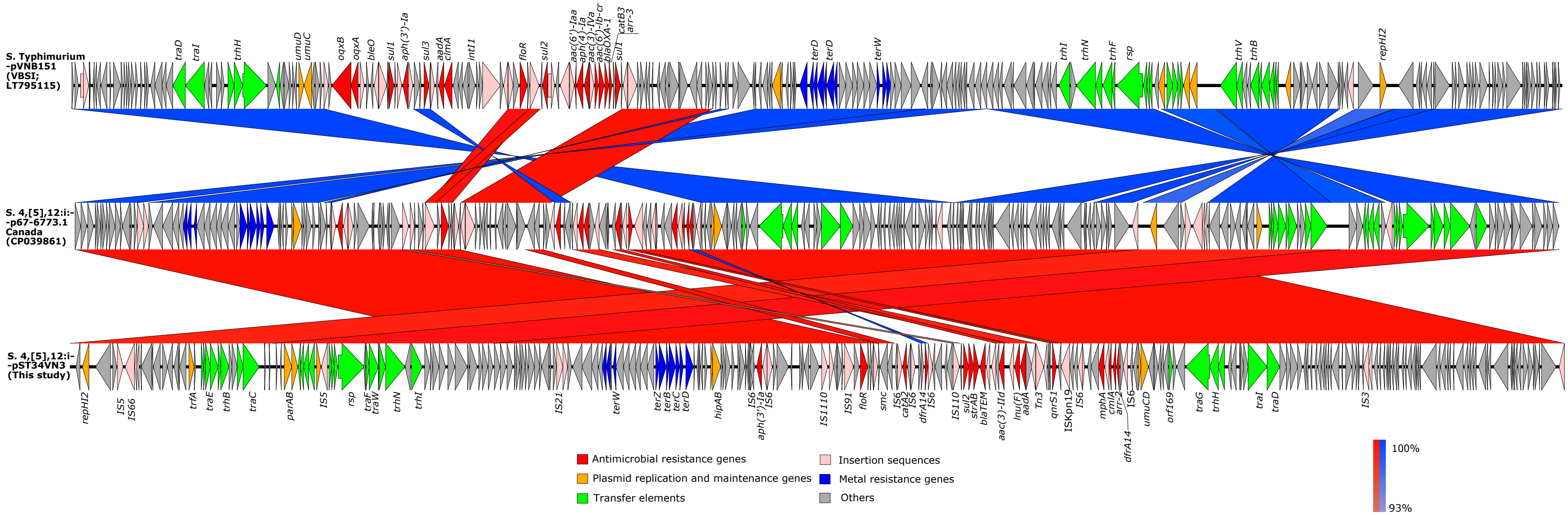


**Figure S4** Comparative genomic of multidrug resistant IncHI2 plasmids recovered from *Salmonella enterica* ST34 (pST34VN3, p67-6773.1, pVNB151). Arrows represent predicted genes (with direction of transcription), which are coloured according to their functions (see keys). The blocks connecting two plasmids indicate regions with high nucleotide similarity (97 – 100%), as either synteny (red) or inversion (blue).

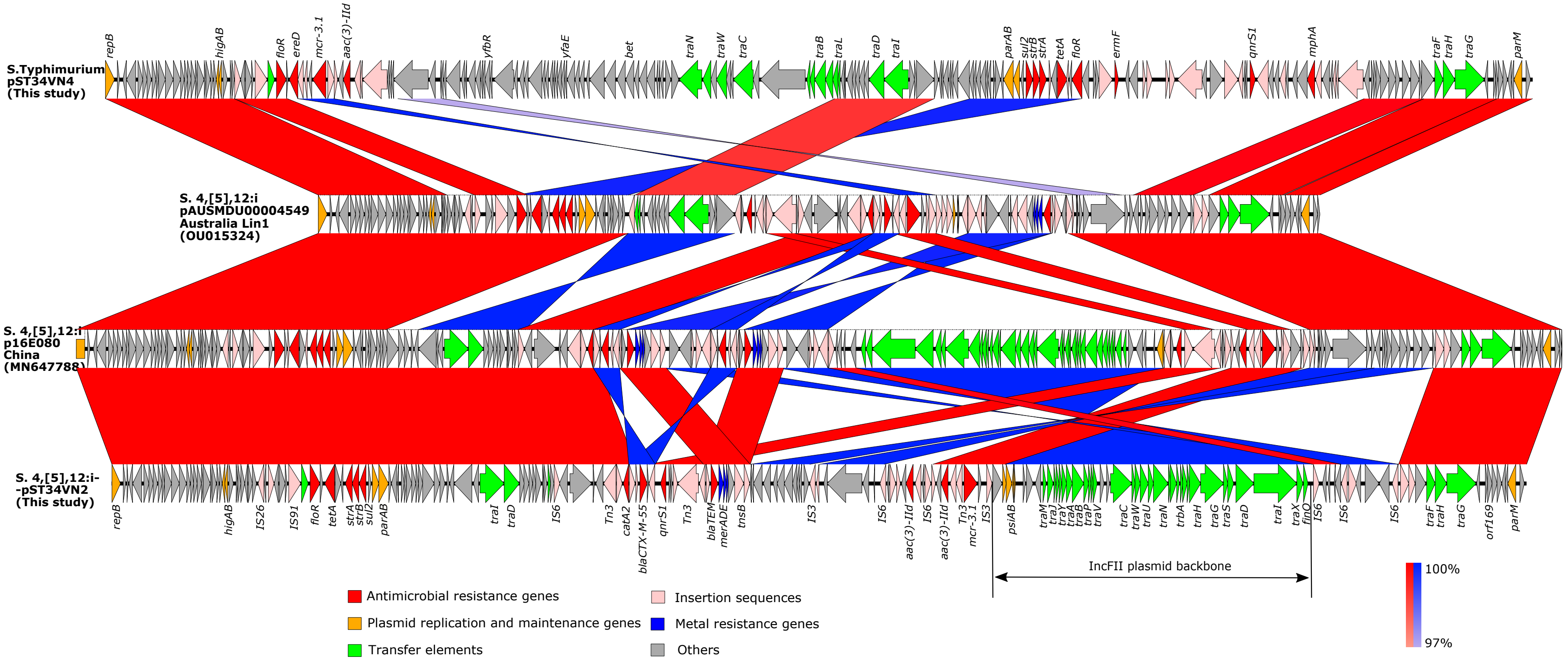


**Figure S5** Comparative genomic of multidrug resistant IncA/C plasmids recovered from *Salmonella enterica* ST34 (pST34VN2, p16E080, pAUSMDU00004549, pST34VN4). Arrows represent predicted genes (with direction of transcription), which are coloured according to their functions (see keys). The blocks connecting two plasmids indicate regions with high nucleotide similarity (97 – 100%), as either synteny (red) or inversion (blue). Bi-directional black arrow delineates the region homologous to the conjugation machinery of the IncFII plasmid (*traM-finO*).

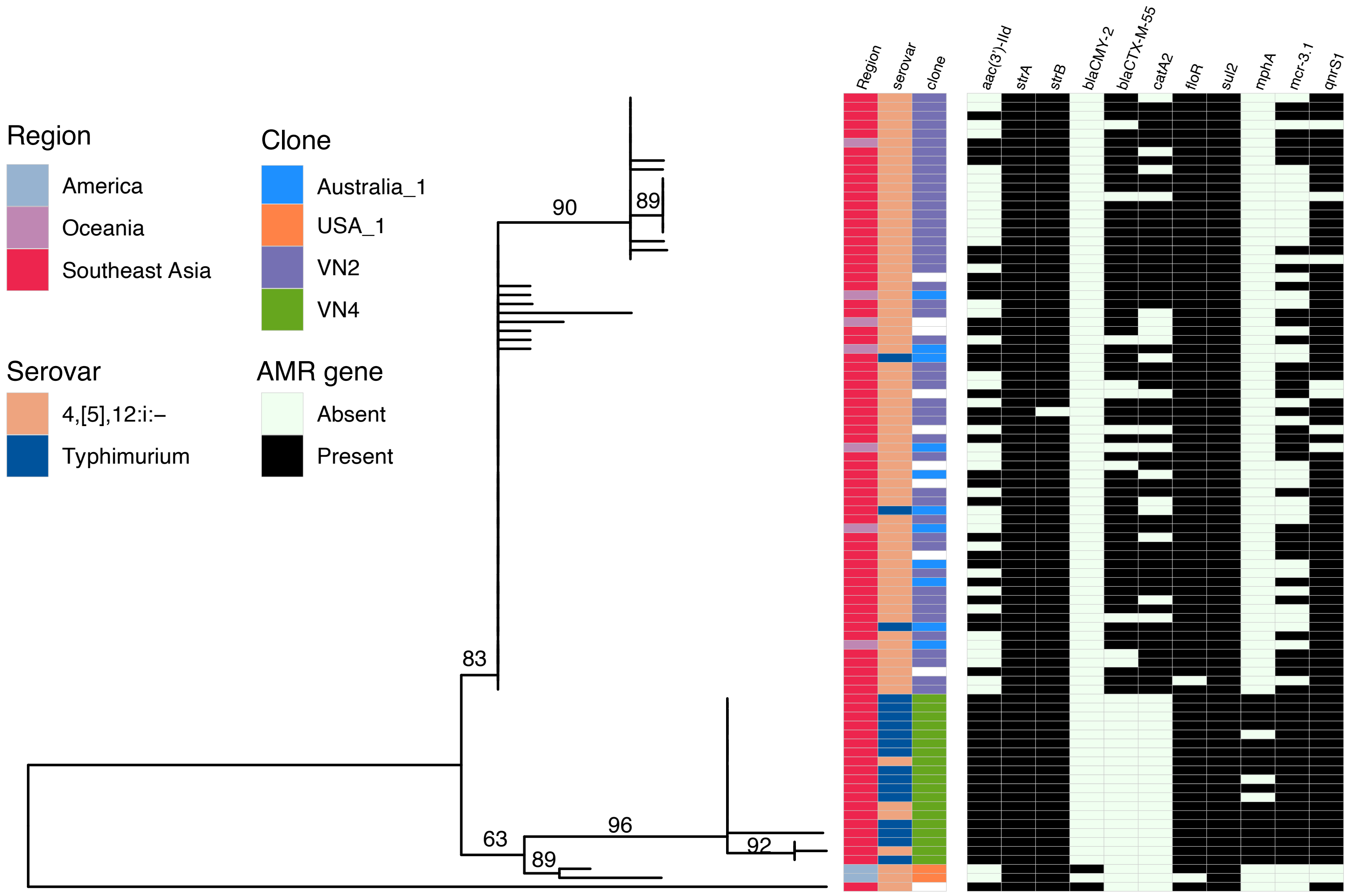


**Figure S6** Plasmid phylogeny of IncA/C2 recovered from *Salmonella enterica* ST34. The figure displays the maximum likelihood phylogeny of IncA/C2 plasmids recovered from 89 ST34 isolates, constructed from 62 single nucleotide polymorphisms. The phylogeny is midpoint rooted. Bootstrap values are displayed at internal branches. The appended heatmap shows data associated with each taxon, including region of origin, predicted serovar, ST34 clones, and presence of AMR genes.

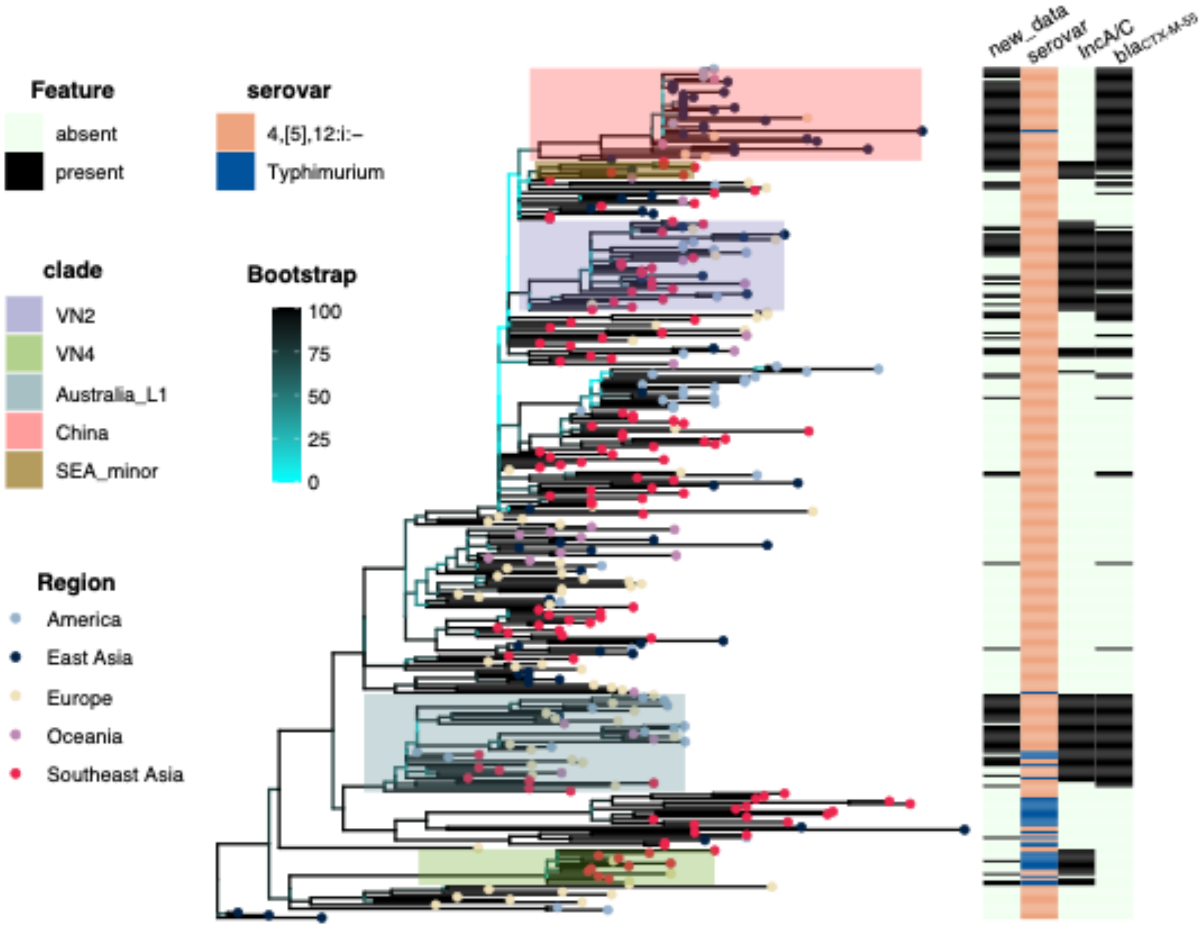


**Figure S7** Global phylogeny of *Salmonella enterica* ST34. The figure displays the maximum likelihood phylogeny of 329 ST34 isolates, combining 222 representative genomes used previously for temporal phylogenetics (Figure 2) and 107 new genomes carrying *bla*$_{CTX-M-55}$ or IncA/C2 plasmid (see Methods). The branches are coloured according to bootstrap values, from low (cyan) to high (black), and tips are coloured based on region of origin. The appended heatmap displays information associated with each taxon, showing whether they are new added data, the predicted serotype, and the presence of IncA/C2 plasmid and *bla*$_{CTX-M-55}$. Coloured shades on the phylogeny highlight defined clusters associated with carriage of *bla*$_{CTX-M-55}$ (see Legend).

**Table S2** Genes showing signal of convergent pseudogenization and gene loss during *Salmonella enterica* ST34 evolution.

| Gene | Locus tag | Length (aa) | Product | Stop codon gain | Frameshift mutation | Gene loss | Total events |
|---|---|---|---|---|---|---|---|
| *ydiV* | B0X74_ 08620 | 237 | anti-FlhC(2)FlhD(4) factor YdiV | 3 | 0 | 0 | 3 |
| *cspC* | B0X74_ 11245 | 69 | RNA chaperone CspC, essential for *phoPQ* regulon in mildly acidic condition | 4 | 6 | 2 | 12 |
| *B0X74_ 16995* | B0X74_ 16995 | 295 | transcriptional regulator | 3 | 1 | 0 | 4 |
| *rpoS* | B0X74_ 17285 | 332 | RNA polymerase sigma factor RpoS | 5 | 16 | 1 | 22 |
| *btuB* | B0X74_ 23885 | 614 | vitamin B12 transporter BtuB | 8 | 5 | 0 | 13 |
